# A *cis*-carotene derived apocarotenoid regulates etioplast and chloroplast development

**DOI:** 10.1101/528331

**Authors:** Christopher I Cazzonelli, Xin Hou, Yagiz Alagoz, John Rivers, Namraj Dhami, Jiwon Lee, Marri Shashikanth, Barry J Pogson

**Affiliations:** Hawkesbury Institute for the Environment, Western Sydney University, Locked Bag 1797, Penrith NSW 2751, Australia.; Australian Research Council Centre of Excellence in Plant Energy Biology, Research School of Biology, The Australian National University, Canberra, ACT 2601, Australia.; Centre for Advanced Microscopy, The Australian National University, Canberra, ACT 2601, Australia

**Author notes:** These two authors contributed equally (Christopher I Cazzonelli, Xin Hou). Address correspondence to and. The author(s) responsible for distribution of materials integral to the findings presented in this article in accordance with the policy described in the Instructions for Authors (www.plantcell.org) are: Barry Pogson and Christopher Cazzonelli.

**Keywords:** carotenoid, post-transcriptional regulation, apocarotenoid signal, prolamellar body, etioplast, photoperiod

## Abstract

Carotenoids are core plastid components, yet a regulatory function during plastid biogenesis remains enigmatic. A unique carotenoid biosynthesis mutant, *carotenoid chloroplast regulation 2* (*ccr2*), that has no prolamellar body (PLB) and normal PROTOCHLOROPHYLLIDE OXIDOREDUCTASE (POR) levels, was used to demonstrate a regulatory function for carotenoids under varied dark-light regimes. A forward genetics approach revealed how an epistatic interaction between a *(-carotene isomerase* mutant (*ziso-155*) and *ccr2* blocked the biosynthesis of specific *cis*-carotenes and restored PLB formation in etioplasts. We attributed this to a novel apocarotenoid signal, as chemical inhibition of carotenoid cleavage dioxygenase activity restored PLB formation in *ccr2* etioplasts during skotomorphogenesis. The apocarotenoid acted in parallel to the transcriptional repressor of photomorphogenesis, DEETIOLATED1 (DET1), to post-transcriptionally regulate PROTOCHLOROPHYLLIDE OXIDOREDUCTASE (POR), PHYTOCHROME INTERACTING FACTOR3 (PIF3) and ELONGATED HYPOCOTYL5 (HY5) protein levels. The apocarotenoid signal and *det1* complemented each other to restore POR levels and PLB formation, thereby controlling plastid development.

**One-sentence summary:** Carotenoids are not just required as core components for plastid biogenesis, they can be cleaved into an apocarotenoid signal that regulates etioplast and chloroplast development during extended periods of darkness.

## INTRODUCTION

Carotenoids are a diverse group of hydrophobic isoprenoid pigments required for numerous biological processes in photosynthetic organisms and are essential for human health (Cazzonelli, 2011; Baranski and Cazzonelli, 2016). In addition to providing plant flowers, fruits and seeds with distinct colours, carotenoids have accessory roles in facilitating the assembly of the light harvesting complex, light capture during photosynthesis and photoprotection during high light and/or temperature stress (Nisar et al., 2015; Baranski and Cazzonelli, 2016). The current frontiers are to discover the regulators of carotenoid biosynthesis, storage, and catabolism and apocarotenoids that in turn regulate plant development and photosynthesis (Cazzonelli and Pogson, 2010; Havaux, 2014; Baranski and Cazzonelli, 2016; Hou et al., 2016).

In higher plants, *cis*-carotene biosynthesis is initiated by the condensation of two molecules of geranylgeranyl diphosphate (GGPP) to form phytoene, which is catalyzed by the rate-limiting enzyme phytoene synthase (PSY) (von Lintig et al., 1997; Li et al., 2008; Rodriguez-Villalon et al., 2009; Welsch et al., 2010; Zhou et al., 2015) (Supplementary Figure 1A). Next, phytoene desaturase (PDS), ζ-carotene desaturases (ZDS), ζ-carotene isomerase (ZISO) and *cis*-*trans*-carotene isomerase (CRTISO) convert the colourless phytoene into the pinkish-red coloured all-*trans*-lycopene (Bartley et al., 1999; Isaacson et al., 2002; Park et al., 2002; Dong et al., 2007; Chen et al., 2010; Yu et al., 2011). In the dark, the isomerisation of tri-*cis*-ζ-carotene to di-*cis*-ζ-carotene and tetra-*cis*-lycopene to all-*trans*-lycopene has a strict requirement for ZISO and CRTISO activity respectively (Park et al., 2002; Chen et al., 2010). However, light-mediated photoisomerisation in the presence of a photosensitiser can substitute for a lack of isomerase activity (Giuliano et al., 2002; Vijayalakshmi et al., 2015; Alagoz et al., 2018).

The carotenoid biosynthetic pathway branches after lycopene to produce α/β-carotenes (Cunningham et al., 1993; Cunningham et al., 1996; Pecker et al., 1996; Ronen et al., 1999). Next, α-carotene and β-carotene are further hydroxylated to produce the oxygenated carotenoids called xanthophylls (e.g. lutein, violaxanthin and zeaxanthin), which comprise the most abundant carotenoids found in photosynthetic leaves. Carotenoids are precursors for apocarotenoids (carotenoid cleavage products) such as phytohormones abscisic acid (ABA) and strigolactone (SL) as well as other apocarotenoids that function in root-mycorrhizal interactions, leaf development, acclimation to environmental stress and retrograde signaling (Havaux, 2014; Walter et al., 2015; Chan et al., 2016; Hou et al., 2016). The carotenoid cleavage dioxygenase and nine-*cis*-epoxy-carotenoid dioxygenase (CCD/NCED) family cleave carotenoids to yield apocarotenoids (Hou et al., 2016). The CCDs have substrate preferences depending on the tissue and nature of the assay (Walter and Strack, 2011; Harrison and Bugg, 2014; Bruno et al., 2016). The five members of the NCED sub-group are exclusively involved in cleavage of violaxanthin and neoxanthin to form ABA (Finkelstein, 2013). The four CCDs have well defined roles in carotenoid degradation in seeds (CCD1 and CCD4) and the synthesis of strigolactones (CCD7/MAX3 and CCD8/MAX4) (Auldridge et al., 2006; Gonzalez-Jorge et al., 2013; Ilg et al., 2014; Al-Babili and Bouwmeester, 2015). Non-enzymatic oxidative cleavage of carotenoids can also generate apocarotenoids by singlet oxygen (^1^O2)-mediated photo-oxidation or by lipoxygenase and peroxidase-mediated co-oxidation (Leenhardt et al., 2006; Gonzalez-Perez et al., 2011). Non-enzymatic carotenoid degradation acts preferentially on selective molecules such as β-carotene and its apocarotenoid derivatives.

*cis*-carotenes such as phytoene, phytofluene and tetra-*cis*-lycopene are reported to be resistant to non-enzymatic degradation (Schaub et al., 2018), although there are some reports that CCDs cleave specific *cis*-carotenes *in vitro* (Bruno et al., 2016). Whether there is a physiological relevance for a *cis*-carotene derived cleavage product or apocarotenoid signal (ACS) *in vivo*, remains unclear. A hunt is on to identify a *cis*-carotene cleavage product that functions as a retrograde signal to regulate nuclear gene expression (Kachanovsky et al., 2012; Fantini et al., 2013; Avendano-Vazquez et al., 2014; Alvarez et al., 2016). CCD4 is implicated in the generation of a *cis*-carotene-derived apocarotenoid signal that regulates leaf shape, chloroplast and nuclear gene expression in the Arabidopsis *clb5/zds* (chloroplast biogenesis-5 / ζ-carotene desaturase) mutant (Avendano-Vazquez et al., 2014). A metabolon regulatory loop around all-*trans*-ζ-carotene was proposed in tomato fruit that can sense *cis*-carotene accumulation, their derivatives or the enzymes themselves (Fantini et al., 2013). The accumulation of *cis*-carotenes in tomato fruit have also been linked to the metabolic feedback-regulation of *PSY* transcription and translation (Kachanovsky et al., 2012; Alvarez et al., 2016). Therefore, *cis*-carotenes themselves or their cleavage products appear to have some functional roles, of which the targets and regulatory mechanism(s) remains unknown.

Determining a mechanistic function for *cis*-carotenes *in planta* has been challenged by low levels of *cis*-carotene accumulation in wild type tissues. Although, when the upper carotenoid pathway is perturbed (Alagoz et al., 2018), seedling lethality (*psy*, *pds* and *zds*), impaired chlorophyll and *cis*-carotene accumulation (*ziso* and *crtiso*) as well as a reduction in lutein (*crtiso*) become apparent (Isaacson et al., 2002; Park et al., 2002). *ziso* mutants in maize (*y9*) and Arabidopsis (*zic*) display pale-green zebra-striping patterns and a delay in cotyledon greening respectively, that resemble a leaf variegation phenotype (Janick-Buckner et al., 2001; Li et al., 2007; Chen et al., 2010). Similarly, *crtiso* loss-of-function in tomato (*tangerine*), melon (*yofi*) and rice (*zebra*) mutants show varying degrees of unexplained yellow leaf variegation (Isaacson et al., 2002; Park et al., 2002; Chai et al., 2010; Galpaz et al., 2013), the causes of which were assumed to relate to perturbed photosystem biogenesis and operation.

During skotomorphogenesis prolamellar bodies (PLB) develop in etioplasts of seedling tissues. The PLB is a crystalline agglomeration of protochlorophyllide (PChlide), POR enzyme and fragments of pro-thylakoid membranes that provide a structural framework for the light-catalysed conversion of PChlide into chlorophylls by POR within picoseconds in conjunction with the assembly of the photosynthetic apparatus (Sundqvist and Dahlin, 1997; Sytina et al., 2008). The de-etiolation of seedlings upon exposure to light activates a sophisticated network consisting of receptors, genetic and biochemical signals that trigger photomorphognesis. Changes in light-induced morphogenesis include: short hypocotyls; expanded and photosynthetically-active cotyledons with developing chloroplasts; and self-regulated stem cell populations at root and shoot apices (Arsovski et al., 2012; Lau and Deng, 2012). Mutants that block skotomorphogenesis and instead promote photomorphogenesis in the dark, such as DETIOLATED1 (DET1) and CONSTITUTIVE PHOTOMORPHOGENIC 1 (COP1) lack POR and thus fail to assemble PLBs (Chory et al., 1989; Sperling et al., 1998; Datta et al., 2006)(Supplementary Figure 1B). This is a consequence of DET1 modulating the levels of PHYTOCHROME INTERACTING FACTOR 3 (PIF3; constitutive transcriptional repressor of photomorphogenesis) and ELONGATED HYPOCOTYL 5 (HY5; positive transcriptional regulator of photomorphogenesis) to control PORA and *PhANG* expression (Stephenson et al., 2009; Lau and Deng, 2012; Xu et al., 2016; Llorente et al., 2017). Thus, in the dark wild-type plants accumulate PIF3, but lack HY5, conversely *det1* lacks PIF3 and accumulates HY5 protein (Supplementary Figure 1B).

PLB formation occurs in carotenoid deficient mutants. Norflurazon treated wheat seedlings grown in darkness can still form a PLB, however it is somewhat aberrant having a looser attachment of POR to the lipid phase and there is an early dissociation from the membranes during photomorphogenesis (Denev et al., 2005). In contrast, *ccr2* is similar to *cop1*/*det1* mutants in that it lacks a PLB in etioplasts, yet it is unique in having normal PChlide and POR protein levels (Park et al., 2002). The associated hyper accumulation of *cis*-carotenes led to the untested hypothesis that *cis*-carotenes structurally prevent PLB formation in etioplasts of dark germinated *ccr2* during skotomorphogenesis and this in turn delayed cotyledon greening following illumination (Park et al., 2002; Datta et al., 2006; Cuttriss et al., 2007). However, it was never apparent why other linear carotenes, such as 15-*cis*-phytoene and all-*trans*-lycopene, permitted PLB formation, raising the question as to whether there were regulatory functions for the *cis*-carotenes that accumulate in *ccr2*.

In this paper, we describe how changes in photoperiod are sufficient to perturb or permit plastid development in *ccr2*, the former leading to leaf variegation. A revertant screen of *ccr2* revealed new connections between a *cis*-carotene-derived signaling metabolite, PLB formation, skotomorphogenesis and chloroplast development. We demonstrate how an unidentified apocarotenoid signal acts in parallel to DET1 to regulate PLB formation and post-transcriptionally control POR, PIF3 and HY5 protein levels in order to fine-tune plastid development in tissues exposed to extended periods of darkness.

## RESULTS

### A shorter photoperiod perturbs chloroplast biogenesis and promotes leaf variegation

The *crtiso* mutants have been reported to display different leaf pigmentation phenotypes (resembling variegations of yellow and green sectors) in a species-dependant manner, with rice and tomato showing changes in pigmentation, but not Arabidopsis. To address if this is species-dependent we investigated if light regimes affected leaf pigment levels and hence plastid development in Arabidopsis *crtiso* mutants. Growing *ccr2* plants at a lower light intensity of 50 E during a long 16 h photoperiod did not cause any obvious changes in morphology or leaf variegation (Supplemental Figure 2A). Whereas, reducing the photoperiod to 8 h resulted in the newly emerged *ccr2* leaves to appear yellow in variegation (Supplemental Figure 2B) due to a substantial reduction in total chlorophyll (Supplemental Figure 2D). As development progressed the yellow leaf (YL) phenotype became less obvious and greener leaves (GL) developed (Supplemental Figure 2C). Therefore, by reducing the photoperiod we were able to replicate in Arabidopsis previous reports in tomato and rice of leaf variegation (Isaacson et al., 2002; Chai et al., 2010).

Next, we demonstrated that day length affects plastid development in newly emerged leaf tissues undergoing cellular differentiation. We replicated the YL phenotype by shifting three weeks old *ccr2* plants from a long 16-h to shorter 8-h photoperiod (Figure 1A-B). The newly emerged leaves of *ccr2* appeared yellow and virescent, while leaves that developed under a 16-h photoperiod remained green similar to wild type (Figure 1B). Consistent with the phenotype, the yellow sectors of *ccr2* displayed a 2.4-fold reduction in total chlorophyll levels, while mature green leaf sectors formed prior to the photoperiod shift had chlorophyll levels similar to that of WT (Figure 1C). The chlorophyll *a*/*b* as well as carotenoid/chlorophyll ratios were not significantly different (Figure 1C). Consistent with the reduction in chlorophyll, total carotenoid content in yellow sectors of *ccr2* was reduced due to lower levels of lutein, β-carotene and neoxanthin (Figure 1D). The percentage composition of zeaxanthin and antheraxanthin was significantly enhanced in yellow sectors, perhaps reflecting a greater demand for xanthophyll cycle pigments that reduce photooxidative damage (Supplemental Figure 2E). Transmission electron microscopy (TEM) revealed that yellow *ccr2* leaf sectors contained poorly differentiated chloroplasts lacking membrane structures consisting of thylakoid and grana stacks, as well as appearing spherical in shape, rather than oval when compared to green leaf tissues from WT or *ccr2* (Figure 1E).

**Figure 1.**
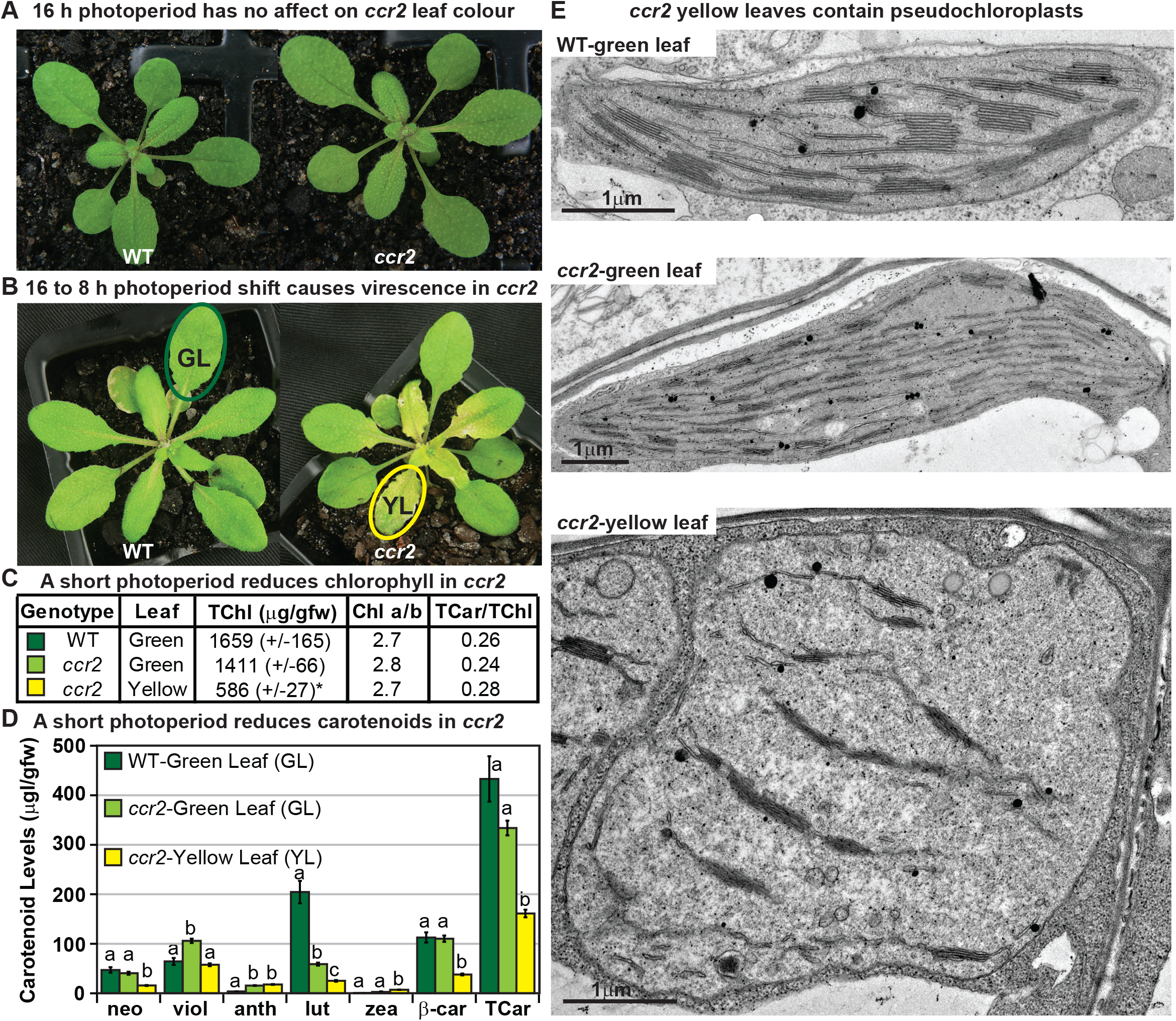
A shorter photoperiod alters plastid development and pigmentation in *ccr2*. **(A)** Three-week-old wild type (WT) and *ccr2* plants growing under a 16-h light photoperiod. **(B)** Three-week-old plants were shifted from a 16-h to 8-h photoperiod for one week and newly emerged or expanded leaves appeared yellow in *ccr2* (YL; yellow outline), while WT displayed green leaves (GL; green outline). **(C)** Chlorophyll levels (μg/gfw) and pigment ratios in green (WT and *ccr2*) and yellow (*ccr2*) leaves formed one week after a photoperiod shift from 16 h to 8 h. Standard error is shown for TChl (n=5, single leaf from 5 plants). Star denotes significant differences (ANOVA; *p* < 0.05). **(D)** Absolute carotenoid levels (μg/gfw) in green (WT and *ccr2*) and yellow (*ccr2*) leaves formed one week after a photoperiod light shift from 16 h to 8 h. Values represent average and standard error bars are displayed (n=5, single leaf from 5 plants). Lettering denotes significance (ANOVA; *p* < 0.05). Neoxanthin (neo), violaxanthin (viol), antheraxanthin (anth), lutein (lutein), zeaxanthin (zea), β-car (β-carotene), Total Chlorophyll (TChl), Chlorophyll a/b ratio (Chl a/b), Total carotenoids (TCar). **(E)** Transmission electron micrograph images showing representative chloroplasts from WT and *ccr2* green leaf sectors as well as yellow leaf sectors of *ccr2*.

**Figure 2.**
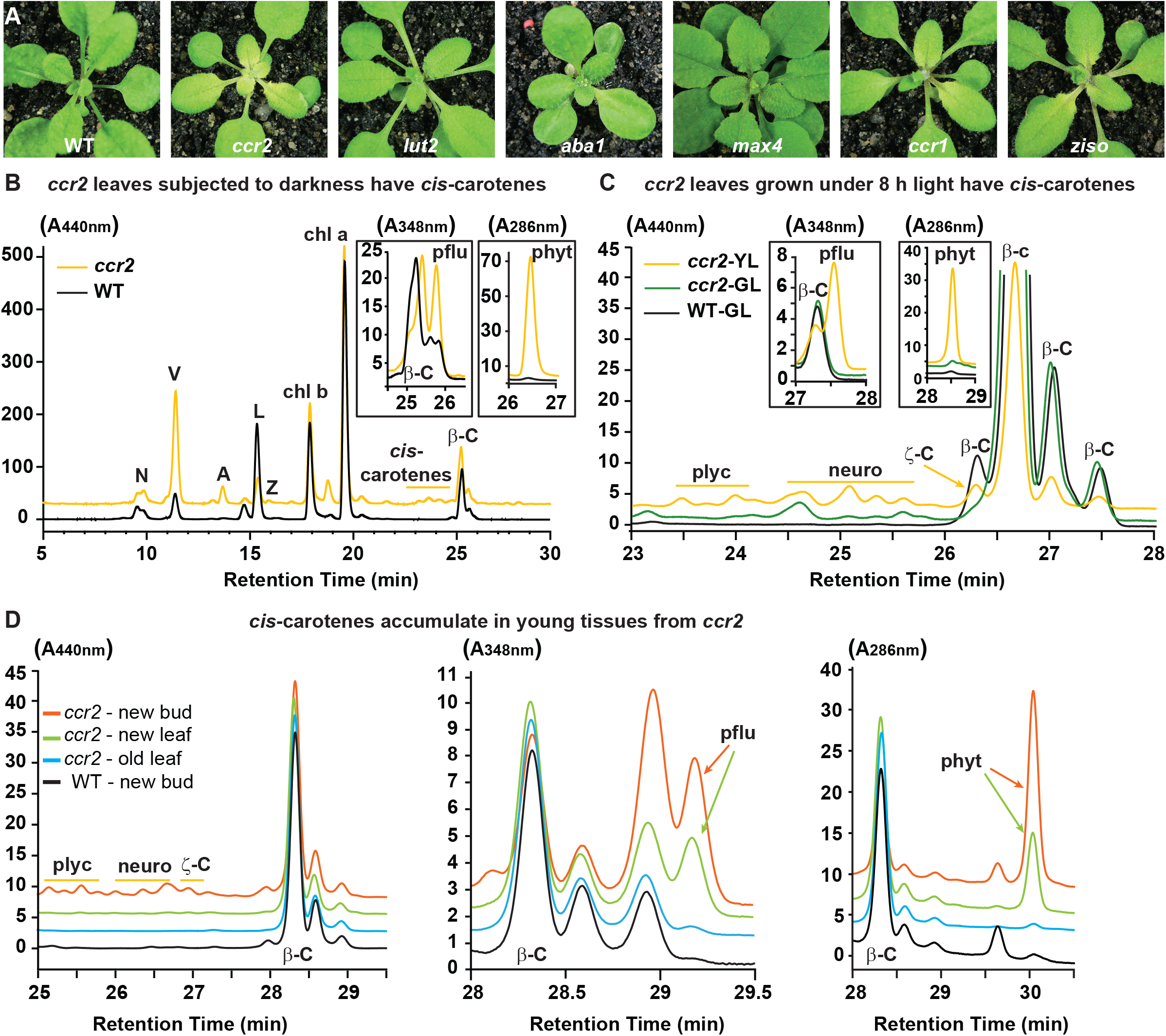
Altered plastid development in *ccr2* is linked with *cis*-carotene accumulation and not to a perturbation in ABA or SL. **(A)** Mutants that perturb the levels of lutein, ABA, SL and accumulate *cis*-carotenes (*ccr2*, *ccr1* and *ziso*) were grown for two weeks under a 16-h photoperiod and then shifted to a shorter 8-h photoperiod for one week. Representative images showing newly emerged and expanding leaves from multiple experimental and biological repetitions (n > 20 plants per line) are displayed. Genetic alleles tested include Col-0 (WT), *ccr2.1* (carotenoid isomerase), *lut2.1* (epsilon lycopene cyclase), *aba1-3* (Ler background) (zeaxanthin epoxidase), *max4/ccd8* (carotenoid cleavage dioxygenase 8), *ccr1.1/sdg8* (set domain group 8) and *ziso1-3* (ζ-carotene isomerase). **(B)** Carotenoid profiles in rosette leaves from three-week-old plants grown under a 16-h photoperiod and subjected to 6-d of extended darkness. **(C)** Carotenoid profiles in three-week-old rosette leaves from plants grown under a constant 8-h light photoperiod. Pigments were profiled in a yellow leaf (YL) and green leaf (GL) from WT and *ccr2*. **(D)** Carotenoid profiles in newly emerged floral bud and rosette leaf tissues harvested from four-week-old plants growing under a 16-h photoperiod. Carotenoid profile traces of various tissue extracts from wild type (WT) and *ccr2* show pigments at wavelengths close to the absorption maxima of A_440nm_ (Neoxanthin; N, violaxanthin; V, antheraxanthin; A, lutein; L, zeaxanthin; Z, β-carotene isomers; β-C, chlorophyll a; Chl a, chlorophyll b; chl b, tetra-*cis-*lycopene; plyc, neurosporene isomers; neuro, and ζ-carotene; ζ-C), A_348nm_ (phytofluene; pflu) and A_286nm_ (phytoene; phyt). HPLC profile y-axis units are in milli-absorbance units (mAU). HPLC traces are representative of multiple leaves from multiple experimental repetitions and retention times vary due to using different columns.

### The leaf variegation phenotype correlated with cis-carotene accumulation

We next investigated the relationship between photoperiod, perturbations in carotenogenesis and plastid development. Green leaf tissues from *ccr2* have an altered proportion of β-xanthophylls at the expense of less lutein, yet plants grown under a longer photoperiod show normal plastid development (Park et al., 2002). This raised a question: does reducing the photoperiod limit the photoisomerisation of tetra-*cis*-lycopene to all-*trans*-lycopene thereby altering lutein, ABA and/or strigolactone biosynthesis? To address this, *ccr2*, *lycopene epsilon cyclase* (*lut2; lutein deficient 2*), *zeaxanthin epoxidase* (*aba1-3; aba deficient 1*) and *carotenoid cleavage dioxygenase 8* (*max 4*; *more axillary branching 4*) mutants were shifted from a 16-h to 8-h photoperiod (Figure 2A). *ccr2* showed a clear yellow variegation phenotype, while the other mutants produced green leaves similar to that of WT. Therefore, we could not attribute the yellow leaf colour variegation to a reduction in lutein or a perturbation of SL or ABA biosynthesis. Next, we tested if the *ccr2* yellow leaf phenotype was linked to the accumulation of *cis*-carotenes in the pathway upstream of all-*trans*-lycopene. Mutations in *PSY, PDS* and *ZDS* cause leaf bleaching and are not viable in soil. Alternatively, *carotenoid chloroplast regulator 1 (ccr1* or otherwise known as *sdg8*; *set domain group 8*) and *(ζ-carotene isomerase* (*ziso*) mutants are viable and accumulate *cis*-carotenes in etiolated tissues (Cazzonelli et al., 2009b; Chen et al., 2010). Indeed, both *ccr1* and *ziso* displayed a partial yellow leaf phenotype near the zone of cellular differentiation (e.g. petiole-leaf margin), however unlike *ccr2* the maturing leaf tissues restore greening rapidly such that *ziso* was more similar to WT than *ccr2* (Figure 2A).

This raised a question: does a shorter photoperiod lead to the accumulation of *cis*-carotenes in newly emerged leaf tissues of *ccr2* displaying altered plastid development? First, we tested if an extended dark period (6 days) would result in the accumulation of *cis*-carotenoids in mature (3 weeks) rosette leaf tissues. Compared to adult WT prolonged darkness resulted in notable yellowing of *ccr2* leaves and clearly discernible accumulation of tetra-*cis*-lycopene, neurosporene isomers, ζ-carotene, phytofluene and phytoene (Figure 2B). We next shifted three-week-old plants from a 16-h to 8-h photoperiod and the yellow sectors from newly emerged *ccr2* leaves accumulated detectable levels of *cis*-lycopene, neurosporene isomers, ζ-carotene, phytofluene and phytoene (Figure 2C). Interestingly, even when plants were grown under a 16-h photoperiod, we could detect phytofluene and phytoene in floral buds as well as newly emerged rosette leaves from *ccr2*, and at trace levels in WT (Figure 2D). In addition, a higher ratio of phytofluene and phytoene relative to β-carotene was observed in newly emerged *ccr2* tissues, which coincided with a lower percentage of lutein when compared to older tissues (Supplemental Table 1).

**Table 1.** A *cis*-carotene derived ACS acts in parallel to DET1 to control PLB formation

| Germplasm | Hypocotyl Length (mm) |  | Apical Hook | Cotyledon | % PLB (-D15) | % PLB (+D15) | *cis*-carotenes |
| --- | --- | --- | --- | --- | --- | --- | --- |
| WT | normal | 13.4 ± 0.2 | yes | closed | 100 | 100 | none detected |
| *ccr2* | normal | 13.8 ± 0.2 | yes | closed | 0 | 85 | phyt, pflu, ζ-C, p-N, p-L |
| *ccr2 det1-154* | shorter | *8.3 ± 0.2 | no | open | 69 | 0 | reduced *cis*-carotenes |
| *det1-154* | shorter | *9.9 ± 0.1 | no | open | ND | ND | phyt, pflu & ζ-C |

ND; not determined; p-N; pro-neurosporene, p-L; pro-lycopene (tetra-*cis*-lycopene), phyt; phytoene, pflu; phytofluene, ζ-c; ζ-carotene, \*; denotes statistical significance (ANOVA, p<0.05).

### Second site genetic reversion restored plastid development in ccr2

We undertook a revertant screen to identify genes and proteins that could complement the plastid development in *ccr2*, while still maintaining a perturbed carotenoid profile. *S*eeds were mutagenized using ethyl-methane sulfonate (EMS), grown and collected from pools of 5-10 M_1_ plants. Approximately 40,000 M_2_ seedlings from 30 stocks of pooled seeds were screened for the emergence of immature green rosette leaves when grown under a 10-h photoperiod. Twenty-five revertant lines reproducibly displayed green immature leaves in response to a photoperiod shift, as exemplified by r*ccr2^154^* and r*ccr2^155^* (Figure 3A). Leaf tissues of all *rccr2* lines contained reduced lutein and xanthophyll composition similar to *ccr2* (Figure 3B). When grown under a shorter photoperiod, *rccr2* lines produced greener rosettes with less yellow colour variegation compared to *ccr2* and chlorophyll levels were similar to WT (Figure 3C-D).

**Figure 3.**
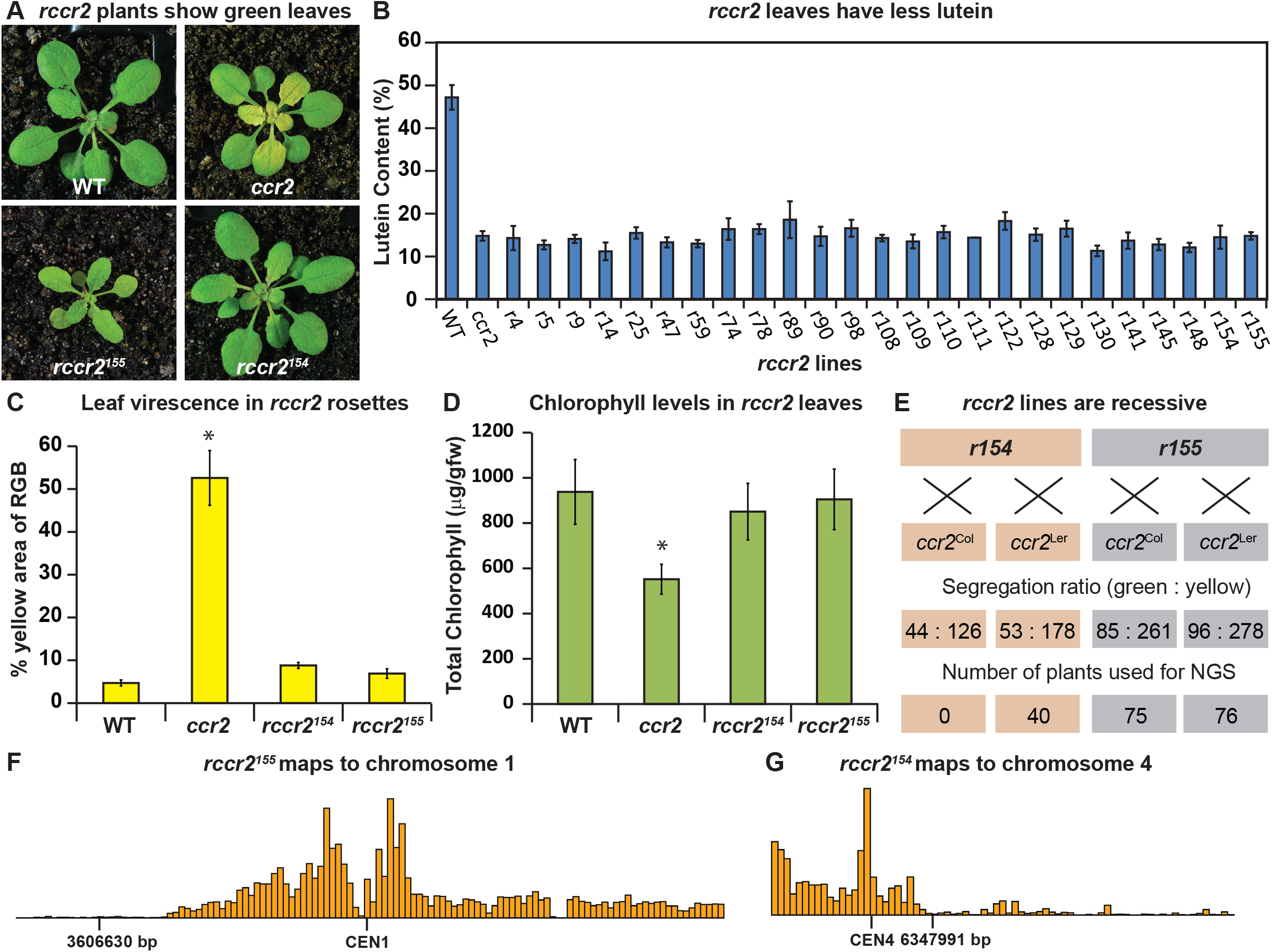
A forward genetics screen identified revertant lines of *ccr2* having reduced lutein and normal chlorophyll accumulation when grown under a shorter photoperiod. **(A)** Representative images of *rccr2^155^* and *rccr2^154^* rosettes one week after shifting two-week old plants from a 16-h to 8-h photoperiod. **(B)** Percentage lutein relative to total carotenoids in immature leaves from WT, *ccr2* and *rccr2* lines. **(C)** The degree of leaf variegation detected in rosettes following a reduction in photoperiod. Leaf variegation (% of yellow relative to RGB) in WT, *ccr2*, *rccr2^154^* and *rccr2^155^* rosettes was quantified using the Lemnatec Scanalyser system and software. **(D)** Total chlorophyll content in rosette leaves from WT, *ccr2*, *rccr2^154^* and *rccr2^155^* plants exposed to a shorter photoperiod. **(E)** Segregation ratios of *rccr2^154^* and *rccr2^155^* after backcrossing to the *ccr2* parent in both Columbia (Col-0) and Landsberg erecta (Ler) ecotypes. (NGS; next generation sequencing) **(F)** and **(G)** NGS of pooled leaf gDNA from a segregating population of *rccr2^155^* **(F)** and *rccr2^154^* **(G)** plants revealed less recombination surrounding SNPs at 3606630 bp and 6347991 bp, respectively. Error bars denote standard error of means (SEM) and stars represent statistical significance (ANOVA; *p* < 0.05).

In order to establish a segregating population for next generation mapping (NGM) *rccr2* lines were backcrossed to the original *ccr2* parent (Col-0) and/or a *ccr2* line established in the Landsberg erecta background (*Lccr2*). All *rccr2* lines were recessive for the reversion of shorter photoperiod dependent yellow leaves (e.g. r*ccr2^154^* and r*ccr2^155^*; Figure 3E). Next generation sequencing (NGS)technologies were used to deep sequence the genomic DNA (gDNA) from leaves of homozygous (M_2_) plants to identify non-recombinant deserts in chromosome 1 (3605576 bp) and chromosome 4 (6346463 bp) for both r*ccr2^155^* and r*ccr2^154^*, respectively (Figure 3F-G). Both non-recombinant deserts contained SNPs displaying a discordant chastity value of approximately 1.0 representing the causal mutation of interest (Austin et al., 2011).

### An epistatic interaction between ziso and ccr2 revealed specific cis-carotenes perturb PLB formation

*rccr2^155^* lacked recombination at the bottom arm of chromosome 1 surrounding a single nucleotide polymorphism (G-A mutation at 3606630 bp) within exon 3 of the *ZISO* gene (639 bp of mRNA), hereafter referred as *ccr2 ziso-155* (Figure 4A). This polymorphism caused a premature stop codon leading to a truncated ZISO protein (212 instead of 367 amino acids). The overexpression of the functional *ZISO* cDNA fragment in *ccr2 ziso-155* restored the leaf variegation phenotype displayed by *ccr2* plants grown under an 8-h photoperiod (Figure 4B). A double mutant generated by crossing *ccr2* with *ziso1-4* further confirmed the loss-of-function in *ziso* can restore plastid development in newly emerged immature leaves of *ccr2*. Carotenoid analysis of immature leaf tissues of *ccr2 ziso-155* revealed reduced lutein and xanthophyll composition similar to *ccr2*, indicating that the complementation of the YL was not due to a change in xanthophyll levels (Figure 3B). The epistatic nature between *ziso* and *crtiso* revealed that a specific *cis*-carotene downstream of *ZISO* activity perturbed plastid development.

Analysis of the *cis*-carotene profile in etiolated cotyledons showed that *ccr2 ziso1-4* had an identical carotenoid profile to that of *ziso* in that it could only accumulate 9,15,9’-tri-*cis*-ζ-carotene, phytofluene and phytoene (Figure 4C). In contrast, *ccr2* accumulated lower levels of these three compounds, yet higher quantities of 9, 9’-di-*cis* ζ-carotene, 7,9,9’-tri-*cis*-neurosporene and 7,9,9’,7’-tetra-*cis*-lycopene, all of which were undetectable in a *ziso* background (Figure 4C). Therefore, *ziso* blocks the biosynthesis of neurosporene isomers, tetra-*cis*-lycopene and 9, 9’-di-*cis* ζ-carotene under shorter photoperiods, and they themselves or their cleavage products appear to disrupt plastid development in *ccr2*.

**Figure 4.**
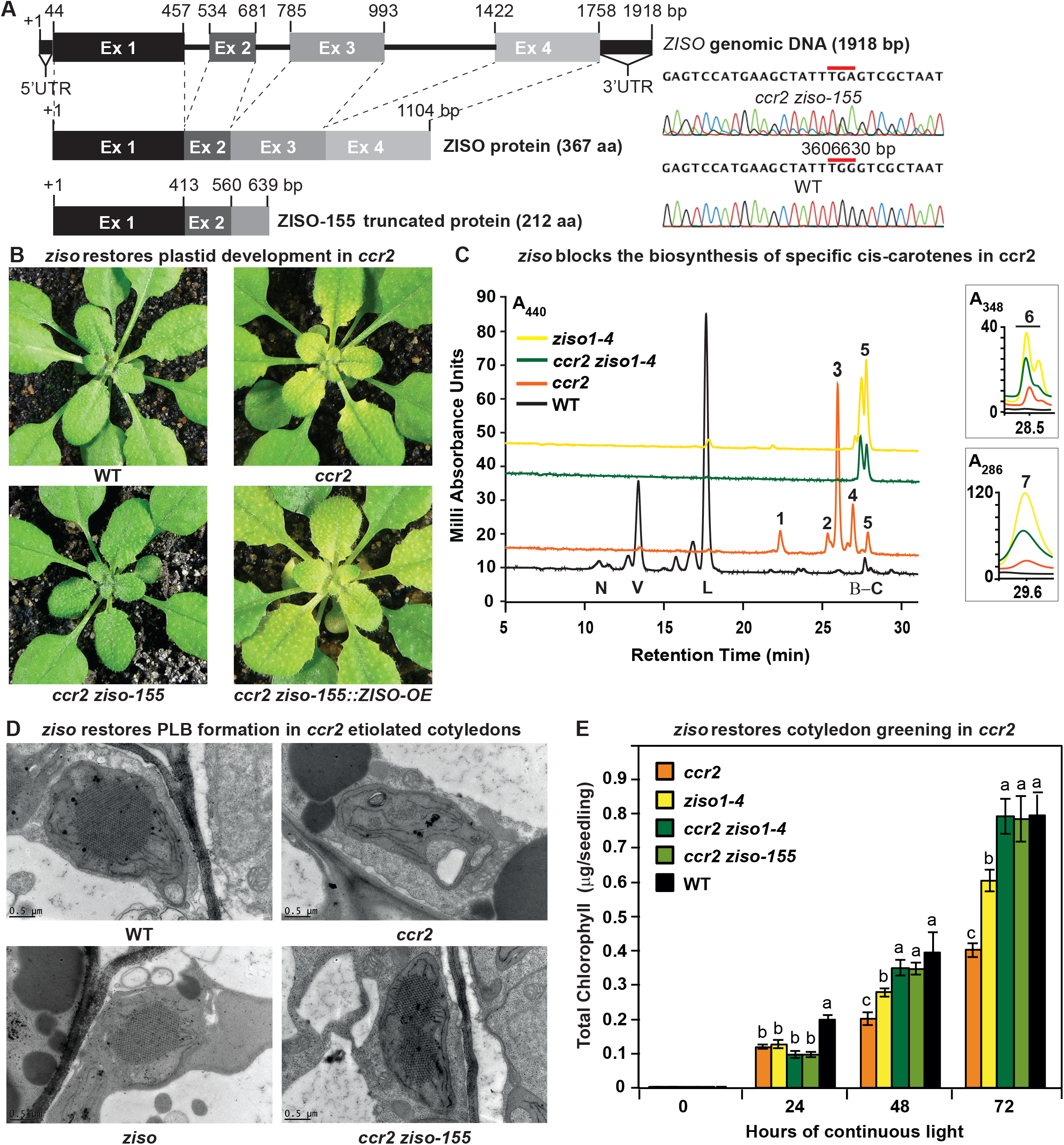
*ziso* alters *cis*-carotene profile to restore PLB formation, plastid development and cotyledon greening in *ccr2*. **(A)** Schematic structure of the wild type *ZISO* gDNA, ZISO protein and the truncated version of the *ZISO-155* genomic sequence. *ccr2 ziso-155* contains a G->A mutation in AT1G10830 (3606630 bp) as confirmed by Sanger sequencing that results in a premature stop codon (TGA) in exon 3. **(B)** Rosette images of WT, *ccr2*, *ccr2 ziso-155*, and *ccr2 ziso-155::ZISO-OE#5* showing leaf pigmentations in newly emerged leaves following a reduction in photoperiod. Images are representative of 84/89 T_4_ generation *ccr2 ziso-155* plants and six independent lines of c*cr2 ziso-155::ZISO-OE*. **(C)** Carotenoid profiles of dark grown cotyledons from WT, *ccr2*, *ziso1-4*, and *ccr2 ziso1-4*. Wavelengths close to the absorption maxima of A_440nm_ (major carotenoids and ζ-carotene isomers), A_348nm_ (phytofluene) and A_286nm_ (phytoene) are shown. Neoxanthin (N); violaxanthin (V); lutein (L); β-carotene (β-C); neurosporene (1 and 2); tetra-*cis*-lycopene (3); pro-neurosporene (4); ζ-carotene (5); phytofluene (6); phytoene (7). **(D)** Transmission electron micrographs of a representative etioplast from 5-d-old dark grown cotyledons. The etioplasts of WT, *ziso* and *ccr2 ziso-155* show well-developed PLBs, while *ccr2* does not have any. Images are representative of 15 plastids from at least 5 TEM sections. **(E)** Total chlorophyll levels in cotyledons following de-etiolation. WT, *ccr2*, *ziso1-4*, *ccr2 ziso-155*, and *ccr2 ziso1-4* were grown in darkness for 4-d, exposed to continuous white light and chlorophyll measured at 0, 24, 48 and 72-h. Letters within a time point denote statistical analysis by ANOVA with a post-hoc Tukey test (n > 20 seedlings). Error bars denote standard error of means (SEM).

How are the specific *cis*-carotenes disrupting plastid development? To answer this question, we first examined etiolated cotyledons of WT, *ccr2*, *ziso* and *ccr2 ziso-155*. We confirmed *ccr2* lacked a PLB in all sections examined (Figure 4D, Supplemental Table 2). We observed 65% of *ziso* etioplasts contained PLBs (Figure 4D, Supplemental Table 2). Intriguingly, the vast majority (>94%) of etioplasts examined from *ccr2 ziso-155* and *ccr2 ziso1-4* contained a PLB (Figure 4D, Supplemental Table 2). Cotyledon greening of de-etiolated seedlings revealed a significant delay in chlorophyll accumulation for both *ccr2* and *ziso1-4* when compared to WT after 24, 48 and 72 h of continuous white light (Figure 4E). The reduced levels of chlorophyll in *ziso* were not as severe as *ccr2*, consistent with *ziso* showing a slight virescent phenotype in comparison to *ccr2* (Figure 2A). Cotyledons of the *ccr2 ziso-155* and *ccr2 ziso1-4* double mutants accumulated levels of chlorophyll similar to that of WT, 48 and 72 h following de-etiolation (Figure 4E). We conclude that a specific *cis*-carotene produced in *ccr2* prevents PLB formation during skotomorphogenesis and perturbs chloroplast development.

### The activation of photosynthesis associated nuclear gene expression restores PLB formation in ccr2

The transcriptomes of WT, *ccr2* and *ccr2 ziso-155* etiolated seedlings (ES), yellow emerging juvenile leaves (JL) from *ccr2*, and green JL leaves from WT and *ccr2 ziso-155* were assessed using RNA sequencing analysis. Compared to WT there were 2-to 4-fold less differentially expressed (DE) genes in *ccr2* (ES;191 and JL;1217) than for *ccr2 ziso-155* (ES;385 and JL;5550). Gene ontology (GO) analysis revealed a DE gene list significantly enriched in metabolic processes and stress responses in both tissue types of *ccr2*. Etiolated tissues of *ccr2* showed DE genes enriched in photosynthetic processes (17/191; FDR < 3.8xE^−06^) that were not apparent in *ccr2 ziso-155*, which had DE genes more responsive to a stimulus (134/382; FDR < 3.7xE^−7^) involving hormones and abiotic stress (Supplemental Table 3). Juvenile leaves of both *ccr2* and *ccr2 ziso-155* showed a significant enrichment in DE genes also responsive to a stimulus (470/1212; FDR < 2.4xE^−34^ and 1724/5510; FDR < 5.4xE^−43^, respectively) involving several hormones and stress. Even more intriguing was the enhanced enrichment of DE genes specific to *ccr2 ziso-155* juvenile leaves that were involved in biological regulation (1623/5510; FDR < 4.2xE^−30^) and epigenetic processes (184/5510; FDR < 3.1xE^−11^) such as DNA methylation, histone modification and gene silencing (Supplemental Table 4).

We utilised Genevestigator to compare DE genes in etiolated seedlings of *ccr2* and *ccr2 ziso-155* with that of mutant germplasm growing on MS media +/-chemical treatments in an attempt to identify co-or contra-changes of gene expression (>20% overlap) (Supplemental Table 3). Norflurazon, a carotenoid inhibitor of PDS activity and inducer of a retrograde signal(s) was able to induce 30-35% of DE genes in *ccr2*, which was not apparent in *ccr2 ziso-155* (12-14%). An unexpected finding was the DE genes in *ccr2* shared 31-42% in common with the *cop9* and *cop1* mutants, which *ccr2 ziso-155* contra-regulated in *cop9*, but not *cop1*. Genes regulated during light-mediated germination were contra-expressed in *ccr2* (28-48%), yet co-expressed in *ccr2 ziso-155* (44-48%).

We next searched for differentially expressed genes in *ccr2* that were attenuated or contra-expressed in the *ccr2 ziso-155*. Twenty contra-expressed genes were identified to be enriched in process related to photosynthesis, pigment biosynthesis and light regulation (5/20; FDR < 1.2xE^−4^) (Supplemental Table 5). Photomorphogenesis associated nuclear gene (*PhMoANG*) expression (e.g. *DET1*, *COP1*) was up-regulated in *ccr2*, yet down-regulated in *ccr2 ziso-155*. This finding is consistent with the fact that DE genes miss-expressed in *ccr2 ziso-155* leaf tissues were enriched in chromatin modifying processes. *det1.1* mutants were shown to have reduced *PIF3* transcripts, and higher *HY5* protein levels that activate downstream *PhANG* expression (Supplemental Table 6)(Lau and Deng, 2012). Indeed, our comparative analysis of contra-expressed genes in *ccr2 ziso-155* revealed the down-regulation of *PIF3*, up-regulation of *HY5* and *PHANG* expression (e.g. *DXS*, *CLB6*, *LHCB1*, *LHCB2*, *RBCS, GUN5*) (Supplemental Table 6). It is not unusual to observe miss-regulation of *PhANG* expression in mutants having impaired plastid development (Ruckle et al., 2007; Woodson et al., 2011). In summary, the repression of negative regulators of photomorphogenesis, correlates well with the up-regulation of *PhANG* expression in *ccr2 ziso-155* and links *cis*-carotene accumulation to gene targets plastid development.

### Activation of photomorphogenesis by det1-154 restores plastid development in ccr2

We searched the SNP deserts of the remaining 24 *rccr2* lines for genes that could link *cis*-carotene signalling to regulators of photomorphogenesis. *rccr2^154^* was mapped to a mutation in de-etiolated 1 (*det1*), hereafter referred as *ccr2 det1-154*, which restored plastid development in immature *ccr2* leaves (Figure 3). Sequencing of the *det1-154* gDNA identified a G to A point mutation at the end of exon 4. Sequencing of the *det1-154* cDNA revealed the removal a 23 amino acid open reading frame due to alternate splicing (Figure 5A). Quantitative PCR analysis confirmed that the shorter *DET1-154* transcript (spliced and missing exon 4) was highly enriched (approx. 200 fold) in *ccr2 det1-154*, while the normal *DET1-154* transcript (contains exon 4) was repressed in *ccr2 det1-154* (Supplemental Figure 3A). The phenotypes of *ccr2 det1-154 and det1-154* were intermediate to that of *det1-1* (Chory et al., 1989) showing a smaller rosette with a shorter floral stem height and reduced fertility relative to the WT (Supplemental Figure 3B). The overexpression of the full length *DET1* transcript (*CaMV35s::DET1-OE)* in *ccr2 det1-154* restored the virescent phenotype in *ccr2* leaves from plants grown under an 8-h photoperiod (Figure 5B). Therefore, alternative splicing of *det1* and removal of exon 4 appeared sufficient to restore plastid development in *ccr2* leaves grown under a shorter photoperiod.

**Figure 5.**
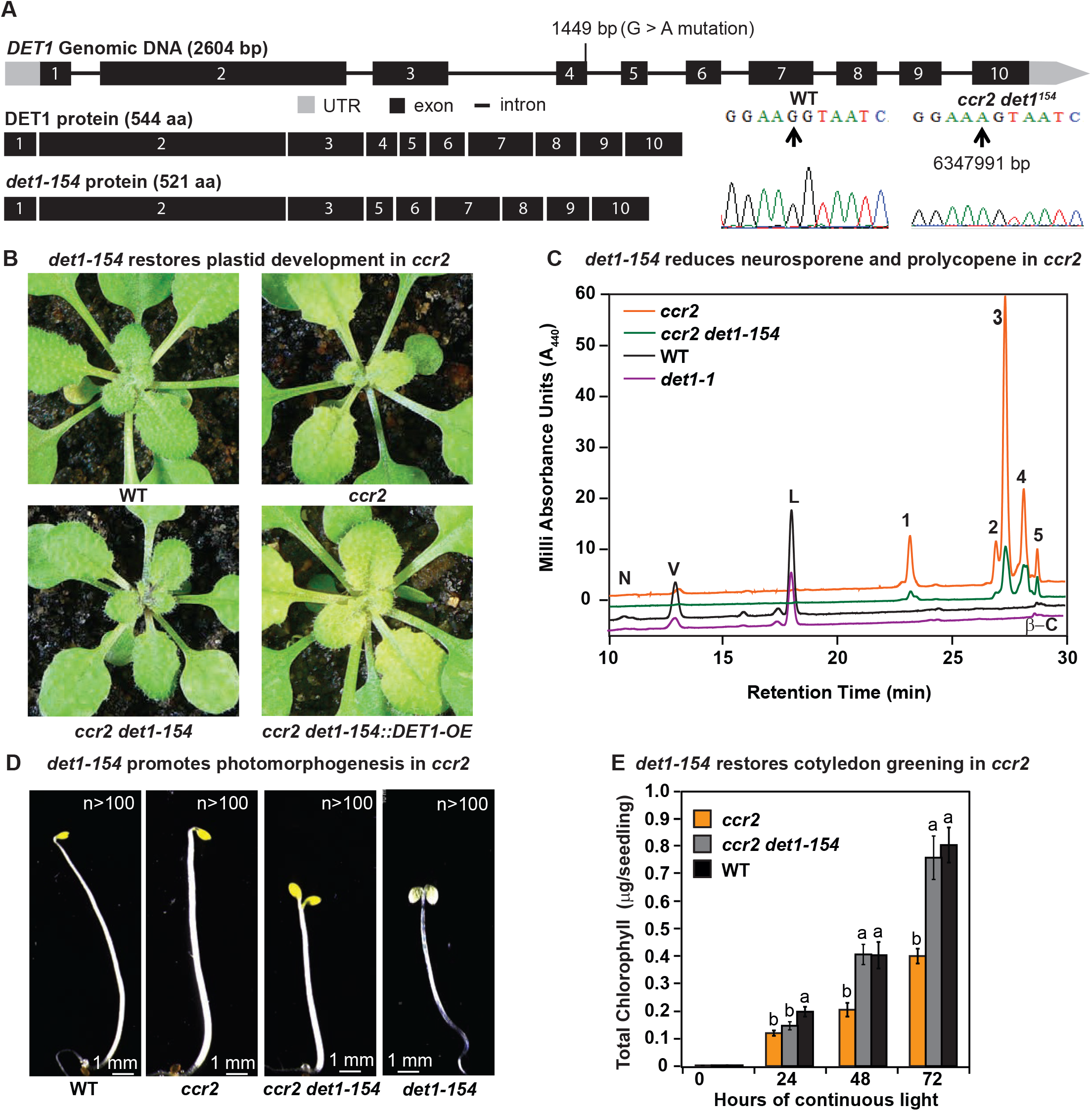
*det1* restores PLB formation, plastid development and cotyledon greening in *ccr2*. **(A)** Schematic structure of the wild type *DET1* gDNA, DET1 protein, SNP confirmation and alternative spliced *DET1-154* protein. A G->A mutation at the end of exon 4 (1449 bp) of AT4G10180 (6347991 bp) was confirmed by Sanger sequencing that leads to the skipping of exon 4 (69 bp). The *DET1-154* splice variant produces a shorter protein (521 aa). Exon 4 comprises 23 amino acids in-frame, having homology to the six-hairpin glycosidase-like (IPR008928) domain. **(B)** Rosette images of WT, *ccr2*, *ccr2 det1-154*, and *ccr2 det1-154::DET1-OE* showing leaf pigmentations in newly emerged leaves following a 16 h to 8 h photoperiod shift assay. Images are representative of 122/149 T_1_ generation *ccr2 det1-154* plants from 12 independent lines surviving Basta herbicide selection after being transformed with pEARLEY::*DET1-OE*. **(C)** Carotenoid profiles of 7-d-old dark grown cotyledons from WT, *ccr2*, *ccr2 det1-154* and *det1-1* etiolated seedlings. Wavelengths close to the absorption maxima of A440 (major carotenoids and ζ-carotene isomers) show neoxanthin (N); violaxanthin (V); lutein (L), β-carotene (β-C) in WT and neurosporene isomers (1 and 2) tetra-*cis*-lycopene (3); pro-neurosporene (4), and pro-ζ-carotene (5) in *ccr2* and to a less extent in *ccr2 det1-154*. **(D)** Etiolated seedling morphology of WT, *ccr2*, *ccr2 det1-154* and *det1-154*. Seedlings were grown in the dark for 7 d on MS media without sucrose. Representative images (>100 seedlings from independent experiments) depict a typical apical hook for WT and *ccr2*, and shorter hypocotyl with open cotyledons for *ccr2 det1-154* and *det1-154*. **(E)** Chlorophyll levels in cotyledons following de-etiolation. *ccr2*, *ccr2 det1-154* and WT were etiolated for 4 d in darkness and thereafter exposed to continuous white light. Chlorophyll measurements were taken at 0 24, 48 and 72 h after de-etiolation. Letters within a time point denote statistical analysis by one-way ANOVA with a post-hoc Tukey test (n > 20 seedlings). Error bars denote standard error of means.

We investigated how *det1-154* can restore plastid development in *ccr2*. *ccr2 det1-154* mature leaves contained less carotenoids and chlorophylls compared to *ccr2* (Supplemental Figure 3C). That is, the xanthophylls and β-carotene were all significantly reduced by *det1-154*. *det1-154* also reduced total *cis*-carotene content in *ccr2* etiolated cotyledons (Figure 5C; Supplemental Figure 3D). However, only tri-*cis*-ζ-carotene, pro-neurosporene and tetra-*cis*-lycopene were significantly reduced in *ccr2 det1-154*, phytoene and phytofluene levels were not significantly different to *ccr2* (; Supplemental Figure 3D). *ccr2* prevented PLB formation during skotomorphogenesis, yet displayed no obvious phenotype that resembled photomorphogenic mutants (Table 1). TEM confirmed that the dark-grown cotyledons from etiolated *ccr2 det1-154* seedlings showed PLBs in 69% of etioplasts examined during skotomorphogenesis (Table 1; Supplemental Figure 3E). The restoration of a PLB in *ccr2 det1-154* dark grown seedlings coincided with a restoration of cotyledon greening following de-etiolation (Figure 5E). In leaves and etiolated cotyledons, *det1* mutants reduced total carotenoid and/or chlorophyll content when compared to WT (Supplemental Table 7). That is, the xanthophylls and β-carotene were all significantly reduced in *det1* mutants. We detected traces of phytoene and phytofluene in emerging leaves and in addition tri-*cis*-ζ-carotene at higher levels in etiolated cotyledons of *det1* mutants (Supplemental Table 7). *det1-154* activated photomorphogenesis in *ccr2* as evident by etiolated seedlings having characteristic shorter hypocotyl, no apical hook and opened large cotyledons similar to *det1-1* (Figure 5D), which agrees with our transcriptomic data whereby *ccr2 det1-154* lead to a repression of *det1* and activation of *PhANGs* (Supplemental Table 6). Therefore, the reduction of the full length *DET1* mRNA in *ccr2* caused a reduction in specific *cis*-carotenes and restored PLB formation (Table 1).

### D15 inhibition of carotenoid cleavage activity reveals a cis-carotene cleavage product controls PLB formation

A question remained as to whether the accumulation of specific *cis*-carotenes lead directly to PLB perturbation as hypothesised (Park et al., 2002), or production of an apocarotenoid signal that could regulate PLB formation. We crossed *ccr2* to carotenoid cleavage dioxygenase loss-of-function mutants; *ccd1*, *ccd4*, *ccd7* (*max3*) and *ccd8* (*max4*) and tested if plants exposed to a shorter photoperiod would revert the virescent leaf phenotype of *ccr2*. We analysed more than 10 plants for each of the *ccr2 ccd* double mutant lines and observed a perturbation in plastid development in >93% of plants, each displaying clearly visible yellow virescent leaves similar to *ccr2* (Supplemental Figure 4A-B). We concluded that no single *ccd* mutant was sufficient to block the production of any *cis*-carotene derived cleavage product. However, there is a degree of functional redundancy among family members, as well as multiple cleavage activities and substrate promiscuity (Hou et al., 2016).

To address this challenge we decided to utilise the aryl-C3N hydroxamic acid compound (D15), which is a specific inhibitor (>70% inhibition) of 9,10 cleavage enzymes (CCD) rather than 11,12 cleavage enzymes (NCED) (Sergeant et al., 2009; Van Norman et al., 2014). We imaged etioplasts from WT and *ccr2* etiolated seedlings treated with an optimal concentration of D15 (Van Norman et al., 2014). The majority (86%) of D15-treated *ccr2* etioplasts displayed a PLB, whilst in control treatments *ccr2* etioplasts showed no discernible PLB (Figure 6A; Supplemental Table 2). Total PChlide levels in WT and *ccr2* after D15 treatment were similar (Figure 6B). As expected, etiolated *ccr2* seedlings grown on D15-treated MS media accumulated chlorophyll in cotyledons within 24-h of continuous light treatment following de-etiolation in a manner similar to WT (Figure 6C). D15 significantly enhanced di-*cis*-ζ-carotene and pro-neurosporene, yet reduced tetra-*cis*-lycopene in etiolated cotyledons of *ccr2* (Figure 6D). In WT etiolated cotyledons, D15 significantly enhanced violaxanthin, neoxanthin and antheraxanthin content, which has previously been shown to also occur in Arabidopsis roots (Van Norman et al., 2014)(Figure 6E). Treatment of dark and light grown wild type seedlings with D15 did not cause adverse pleiotropic effects on germination, hypocotyl elongation and/or plastid development in cotyledons (Figure 6, Table 1, Supplemental Table 2). Therefore, apocarotenoid formation from either cleavage of di-*cis*-ζ-carotene and/or pro-neurosporene in *ccr2* can perturb PLB formation independent of PChlide biosynthesis.

**Figure 6.**
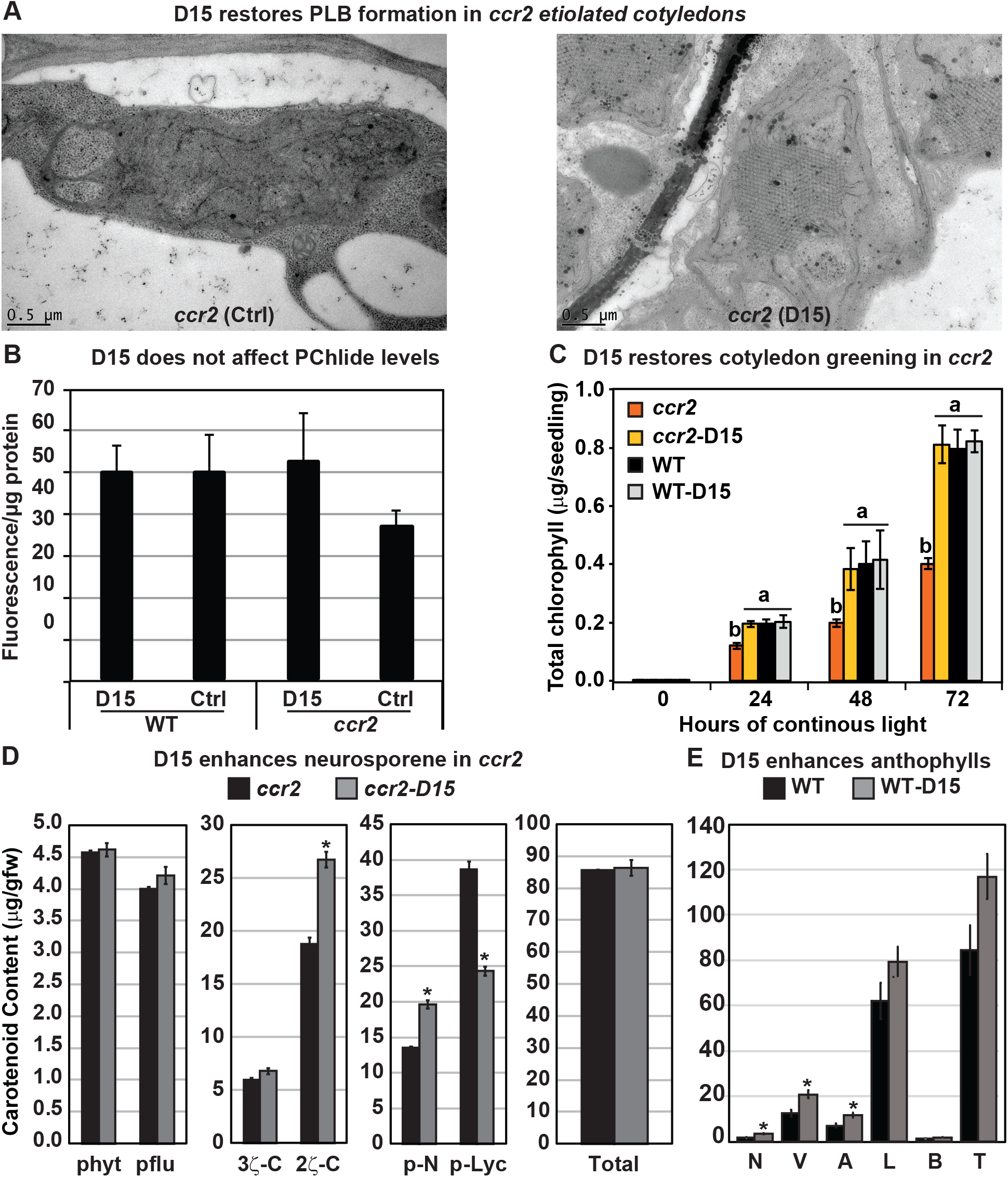
The carotenoid cleavage dioxygenase (CCD) inhibitor, D15, restores PLB formation in etiolated *ccr2* seedlings, cotyledon greening following de-etiolation and alters *cis*-carotene accumulation. **(A)** Transmission electron micrographs of a representative etioplast from 5-d-old dark grown cotyledons reveal a well-developed PLB in *ccr2* treated with the D15, but not in *ccr2* treated with ethanol only (control; ctrl). **(B)** Pchlide levels in Wild Type (WT) and *ccr2* treated +/-D15. Fluorescence was measured at 638 nm and 675 nm with an excitation at 440 nm. Net fluorescence of Pchlide was calculated and normalised to protein content. **(C)** D15 restores chlorophyll accumulation in *ccr2* de-etiolated seedlings exposed to continuous light. Twenty seedlings from each of three biological replicates were harvested for chlorophyll determination in every 24 h under continuous light. Statistical analysis was by ANOVA with a post-hoc Tukey test (n= 20 seedlings). **(D)** *cis*-carotene quantification in etiolated cotyledons of *ccr2* treated with D15. phytoene (phyt), phytofluene (pflu), tri-*cis*-ζ-carotene (3ζ-C), di-*cis*-ζ-carotene (2ζ-C), pro-neurosporene (p-N), tetra-*cis*-lycopene (p-lyc) and total *cis*-carotenes were quantified at absorption wavelengths providing maximum detection. Star denotes significance (ANOVA, *p* < 0.05). Error bars show standard error (n =4). **(E)** Quantification of carotenoid levels in etiolated tissues of WT treated with D15. Neoxanthin (N); violaxanthin (V); antheraxanthin (A), lutein (L), β-carotene (β-C) and total carotenoids (T) were quantified at a 440nm absorption wavelength providing maximum detection. Star denotes significance (ANOVA, *p* < 0.05). Data is representative of two independent experiments.

### A cis-carotene cleavage product acts downstream of DET1 to post-transcriptionally regulate protein levels

We searched for a transcriptional regulatory mechanism by which a *cis*-carotene cleavage product could control PLB formation during skotomorphogenesis. *PORA* transcript levels are relatively high in etiolated seedlings, becoming down-regulated upon exposure to white light or when photomorphogenesis is activated by *det1-1* (Armstrong et al., 1995; Sperling et al., 1998). PIF3 and HY5 are key regulatory transcription factors involved in controlling downstream *PhANG* expression during the dark to light transition (Osterlund et al., 2000; Dong et al., 2014). A reduction in *PORA* and *PIF3* has been shown to perturb PLB formation. In etiolated tissues of *ccr2 det1-154*, which harbour etioplasts containing a PLB, the transcript levels of *PORA, PORB, PIF3* and *HY5* were substantially reduced. D15 treatment did not affect the expression levels of these genes in either WT or *ccr2 det1-154* (Figure 7A), therefore, *cis*-carotene cleavage does not appear to directly affect the transcriptional regulation of these genes.

**Figure 7.**
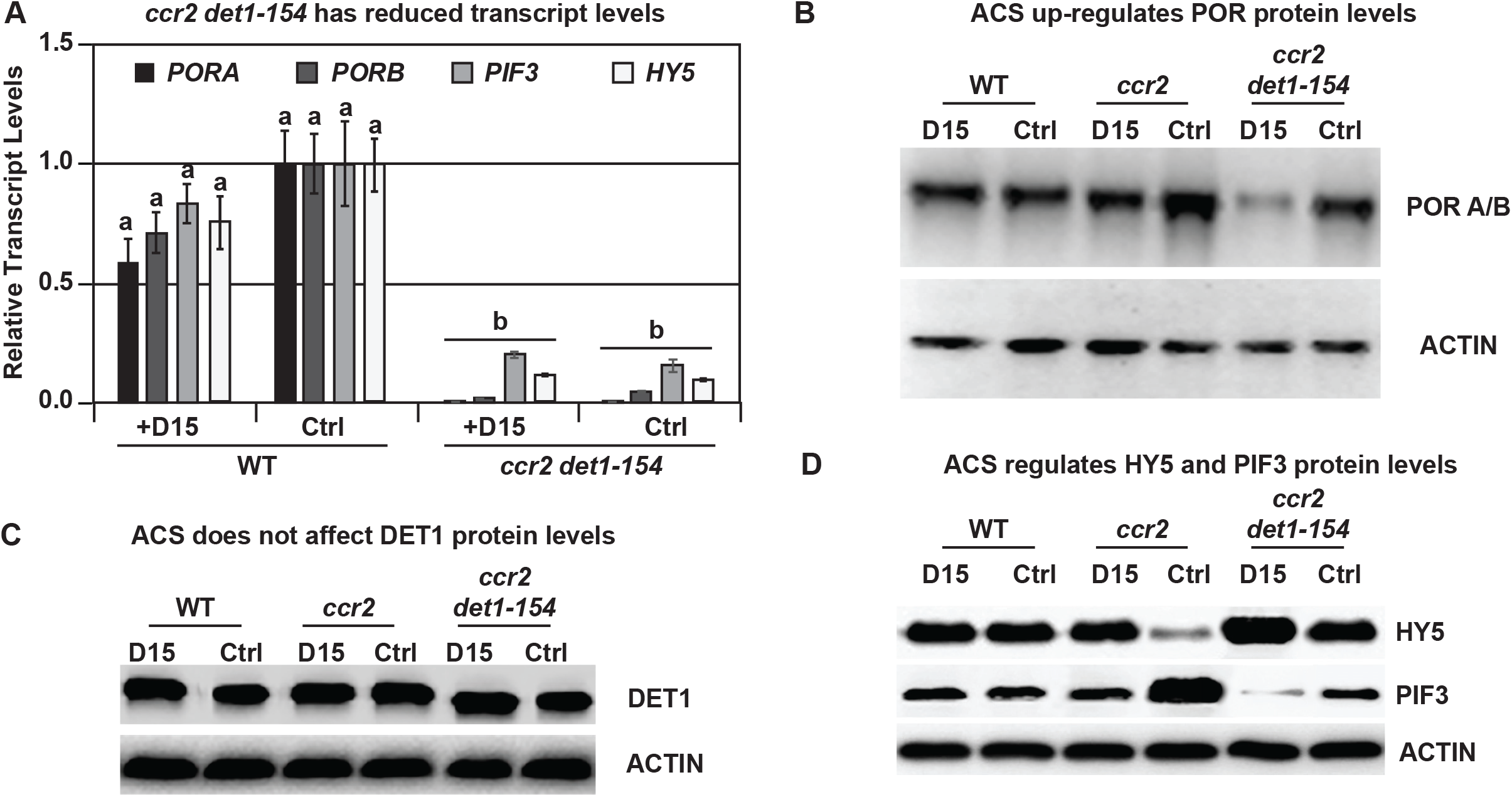
Chemical inhibition of CCD activity identifies a *ccr2* generated cleavage product acts in parallel *det1-154* to post-transcriptionally regulate POR, HY5 and PIF3 protein levels. **(A)** Transcript levels of *PORA*, *PORB, PIF3* and *HY5* in WT and *ccr2 det1-154* etiolated seedlings growing on MS media +/-D15. Statistical analysis was performed using two-way ANOVA followed by a post-hoc paired *t*-test (*p* < 0.05). Error bars represent standard error of means. **(B)** Representative image of a western blot showing POR protein levels. Proteins were extracted from WT, *ccr2* and *ccr2 det1-154* etiolated seedlings grown on MS media +/-D15 (control; Ctrl). The membrane was re-probed using anti-Actin antibody as an internal control to normalise POR protein levels among different samples. **(C)** Representative image of a western blot showing DET1 protein levels. Proteins were extracted from WT, *ccr2* and *ccr2 det1-154* etiolated seedlings grown on MS media +/-D15 (control; Ctrl). The membrane was re-probed using anti-Actin antibody as an internal control to normalise POR protein levels among different samples. **(D)** Western blot image of HY5 and PIF3 protein levels. Proteins were extracted from WT, *ccr2* and *ccr2 det1-154* etiolated seedlings grown on MS media (Ctrl) or media containing D15. The membrane was re-probed using anti-Actin antibody as an internal control to normalise HY5/PIF3 protein levels among different samples.

We then searched for a change in POR protein in dark grown seedlings, with or without D15, noting that wild-type and *ccr2* accumulated POR (Park et al., 2002) and *det1* lacked POR (Sperling et al., 1998)(Supplementary Figure 1B). Under the electrophoresis conditions used herein, the Arabidopsis PORA/B proteins were detected as a single immunoreactive signal (PORA; 37 kDa, and PORB; 36 kD) (Sperling et al., 1998; Park et al., 2002; Paddock et al., 2012) (Figure 7B). A substantial increase in POR was observed in *ccr2*, which was reduced back to WT levels by D15 (Figure 7B). *ccr2 det1-154* accumulated wild-type levels of POR (Figure 7B), complementing the reported lack of POR in etiolated *det1* tissues (Sperling et al., 1998)(Supplementary Figure 1B). Intriguingly, treatment of *ccr2 det1-154* with D15 reverted POR levels back to those expected for *det1*. This was not due to *ccr2* or D15 changing DET1 protein levels (Figure 7C). Therefore, *cis*-carotene cleavage mediates a signal that elevates POR accumulation in *det1*.

DET1 is a negative regulator of photomorphogenesis, such that *det1* mutants lack PIF3 and accumulate HY5 protein levels during skotomorphogenesis (Osterlund et al., 2000; Dong et al., 2014)(Supplementary Figure 1B). We questioned if the apocarotenoid signal acted upstream of the PIF3-HY5 regulatory hub that controls *PhANG* expression, noting that wild-type has high levels of PIF3 and low or trace levels of HY5, with the converse in *det1* (Dong et al., 2014) (Supplementary Figure 1B). PIF3 levels increased and HY5 decreased in both *ccr2* and *ccr2 det1-154* etiolated cotyledons and this was reverted by D15 treatment (Figure 7D). This indicates that an apocarotenoid signal can post-transcriptionally change the PIF3 / HY5 ratio in the presence or absence of DET1, indicating it is acting either in parallel with, or downstream of, DET1. The relative difference in PIF3 levels in *ccr2* compared to *ccr2 det1-154* in the presence of D15 would suggest the two pathways operate in parallel.

## DISCUSSION

Plastid and light signalling coordinate leaf development under various photoperiods, and younger leaves display a greater plasticity to modulate their pigment levels in response to environmental change (Lepisto and Rintamaki, 2012; Dhami et al., 2018). We attribute *ccr2* leaf variegation to the fine-tuning of plastid development in meristematic cells as a consequence of *cis*-carotene accumulation and not the generation of singlet oxygen (Kato et al., 2009; Chai et al., 2010; Han et al., 2012). Our evidence revealed that leaf variegation is linked to the hyper-accumulation of specific *cis*-carotenes since, *ziso-155* and *det1-154* as well as D15 were able to reduce *cis*-carotene biosynthesis in *ccr2* tissues, as well as restore leaf greening in plants grown under a shorter photoperiod (Figures 4,5). A shorter photoperiod maybe a seasonal factor capable of triggering *cis*-carotene hyper-accumulation in newly emerged photosynthetic tissues when CRTISO activity is perturbed, and cause leaf variegation. The altered plastid development in etiolated cotyledons and younger virescent leaves from *ccr2* cannot be attributed to a block in lutein, strigolactone, ABA or alteration in xanthophyll composition (Figure 2). Phytoene, phytofluene and to a lesser extent ζ-carotene were noted to accumulate in wild type tissues from different plant species (Alagoz et al., 2018). We also detected traces of these *cis*-carotenes in newly emerged leaves from wild type, and even more so in *det1* mutants. Without the signal itself to assess the physiological function in wild-type plant tissues, we provided evidence for the existence of a *cis*-carotene cleavage product in *ccr2* that can regulate PLB formation during skotomorphogenesis and plastid development during leaf greening independent of, and capable of compensating for mutations in DET1 (Figure 8A). We contrast how the *cis*-carotene derived novel apocarotenoid signal, in parallel with DET1 can post-transcriptionally control repressor and activator proteins that mediate the expression of a similar set of *PhANGs* in both the etioplast and chloroplast.

**Figure 8.**
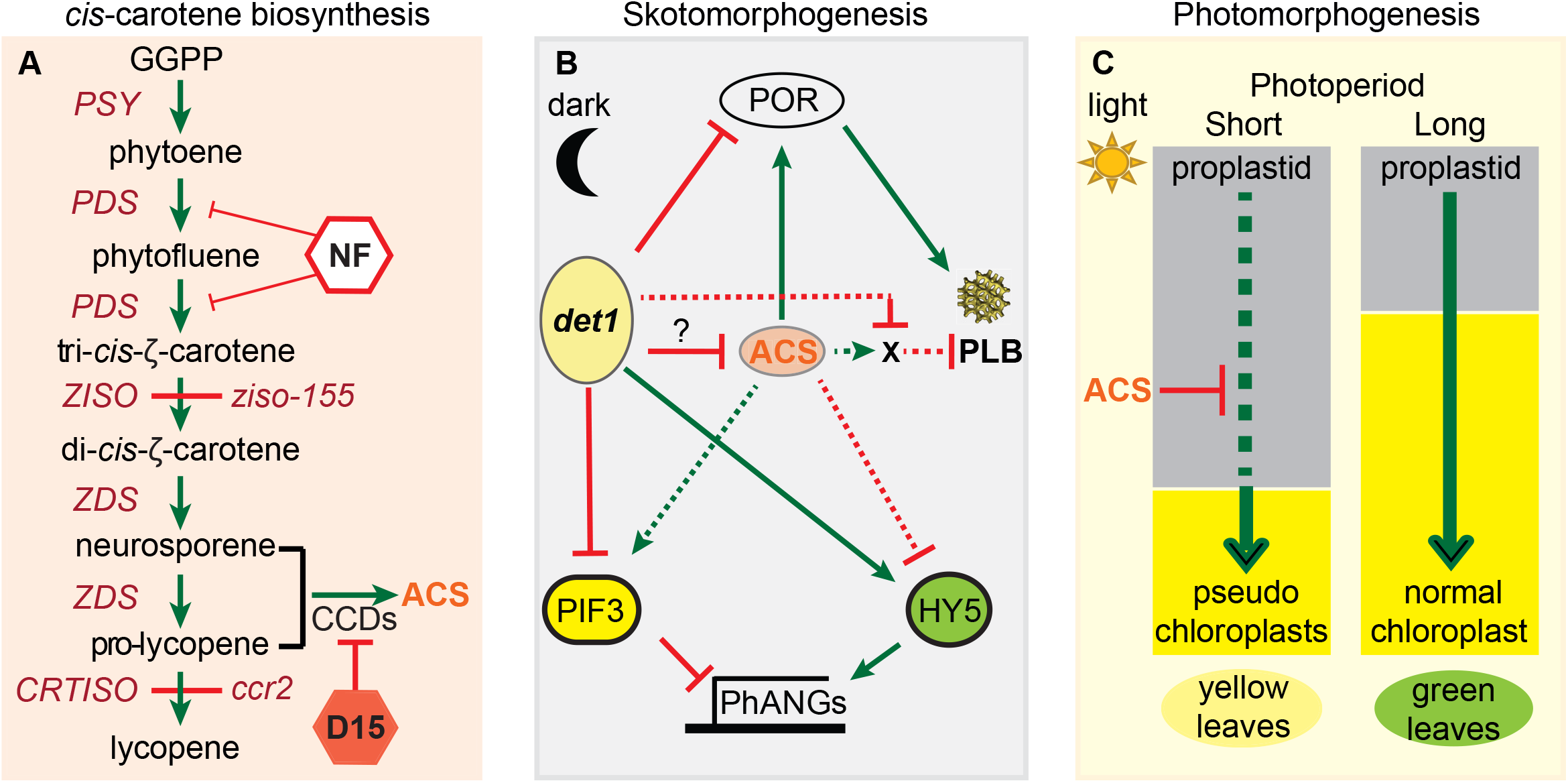
A model describing how a *cis*-carotene derived cleavage product controls POR and PLB formation during plastid development. **(A)** *ccr2* can accumulate poly-*cis*-carotenes that undergo enzymatic cleavage via CCDs to generate an apocarotenoid signal (ACS). Norflurazon (NF) treatment of *ccr2* etiolated seedlings or the loss-of-function in *ziso-155* block the accumulation of downstream cis-carotenes required for the biosynthesis of ACS. Chemical treatment of etiolated seedlings with D15 inhibits CCD cleavage of pro-neurosporene and/or tetra-*cis*-lycopene isomers into ACS. **(B)** During skotomorphogenesis, ACS promotes “Factor X”. Factor X negatively affects PLB formation. Factor X could act to stabilise proteins by disrupting ubiquitination, de-ubiquitination, protease mediated protein degradation, heterodimerization of transcription factors, coactivator concentrations, and/or interact with ligand binding sites of receptors. DET1 is a repressor of photomorphogenesis that post-transcriptionally regulate PIF3 and HY5 protein levels, which control *PhANG* expression. *det1* mutants lack POR and cannot make a PLB. ACS post-transcriptionally enhances POR protein levels, while *det1* blocks Factor X, thereby allowing PLB formation in *ccr2 det1-154*. *det1* reduces *cis*-carotene accumulation, and downregulates pro-neurosporene and tetra-*cis*-lycopene to maintain a threshold level of ACS. **(C)** During photomorphogenesis, extended dark and/or shorter photoperiods, ACS manifests in newly emerged leaves from the *ccr2* shoot meristem and perturbed chloroplast development and chlorophyll accumulation causing a leaf variegation phenotype. Green arrows and red lines represent positive and negative regulation, respectively. Abbreviations: PSY, phytoene synthase; PDS, phytoene desaturase, ZDS, ζ-carotene desaturase; Z1SO, ζ-carotene isomerase; CRTISO, carotenoid isomerase; *det1-154*, DEETIOLATED1-154; D15, inhibitor of CCD activity; CCD, carotenoid cleavage dioxygenase; *ccr2*, *CRTISO* mutant.

### A cis-carotene derived cleavage product regulates plastid development

Due to their hydrophobicity and *cis*-configuration, *cis*-carotenes were proposed to function as a membrane-bound structural inhibitor of PLB formation during skotomorphogenesis (Park et al., 2002; Cuttriss et al., 2007). Instead, we demonstrate here that *ccr2* generated a *cis*-carotene-derived cleavage product, as D15 chemical inhibition of CCD activity (Figure 8A) restored PLB formation (85%) in *ccr2* etioplasts (Figure 6). This is in agreeance with evidence showing *cis*-carotenes are cleavable *in vitro* by CCD7 enzymatic activity (Bruno et al., 2016) and that CCD4 activity is necessary for generation of a *cis*-carotene derived apocarotenoid signal in *zds/clb5*, that affected leaf development (Avendano-Vazquez et al., 2014). However, loss-in-function of *ccd1, ccd4, ccd7* and *ccd8* were not sufficient to restore plastid development and prevent leaf variegation in *ccr2* (Supplemental Figure 4). So, we conclude that there must be some redundancy among two or more CCDs in generating a *ccr2* derived apocarotenoid signalling metabolite that controls plastid development.

Which *cis*-carotene is the precursor for the apocarotenoid signal? Treatment with NF (Figure 8A) restored PLB formation in *ccr2* etioplasts (Cuttriss et al., 2007), which ruled out both phytoene and phytofluene as substrates for the generation of a cleavage product. Here we show *ZISO* restored PLB formation and cotyledon greening in *ccr2* ruling out tri-*cis*-ζ-carotene and revealing that di-*cis*-ζ-carotene, pro-neurosporene isomers and/or tetra-*cis*-lycopene are likely candidates (Figure 4). *ccr2 det1-154* displayed a substantial reduction in pro-neurosporene and tetra-*cis*-lycopene, and to a lesser extent di-*cis* ζ-carotene (Supplemental Figure 3). Tetra-*cis*-lycopene accumulated in variegated leaves from the rice *zebra* mutant (Han et al., 2012). However, in the presence of D15 and hence absence of any enzymatic cleavage, only di-*cis*-ζ-carotene and pro-neurosporene accumulated, not tetra-*cis*-lycopene (Figure 6). Based on the evidence to date, we consider pro-neurosporene and perhaps di-*cis*-ζ-carotene are preferred substrate(s) for *in vivo* cleavage into a signaling metabolite.

### A cis-carotene cleavage product controls PLB formation independent of GUN activity

One question is whether the proposed apocarotenoid requires GUN activity to regulate PLB formation and/or *PhANG* expression? Given that *gun1* etioplasts contain PLBs, then that aspect of the *ccr2* phenotype is not GUN-related (Susek et al., 1993; Xu et al., 2016). Additionally, there were relatively few differentially expressed genes in common between *ccr2* etiolated seedlings and *gun1/gun5* seedlings treated with norflurazon (Supplemental Table 6) and none of the 25 revertants were in genic regions to which *GUN* genes are located. Norflurazon treatment of etiolated tissues does not affect PLB formation in wild type, but can restore PLB formation in *ccr2* (Cuttriss et al., 2007; Xu et al., 2016). Lincomycin treatment, on the other hand can suppress PLB formation in etiolated seedlings and unlike norflurazon, affects the phenotype of *pifq* mutant seedlings grown in the dark. GUN1-facilitated retrograde-signaling antagonized *PIF*-regulated gene expression and attenuated de-etiolation phenotypes triggered by lincomycin (Martin et al., 2016). Furthermore, lincomycin also inhibited PLB formation in the *pifq* mutant, revealing that PIFs are not necessary for PLB formation (Martin et al., 2016). GUN1-dependent and independent signaling pathways were proposed to act upstream of HY5 to repress photomorphogenesis of cotyledons (Ruckle et al., 2007). Intriguingly, the *ccr2* generated *cis*-carotene derived cleavage product also regulated a distinct set of genes involved in a photomorphogenic-dependent pathway. The nature by which a *cis*-carotene derived cleavage product regulates PLB formation by post-transcriptionally enhancing POR is quite distinct to that of GUN regulation of *PhANGs*.

### An apocarotenoid post-transcriptionally regulates PIF3 and HY5 protein levels

Here we demonstrate that the *ccr2*-generated apocarotenoid acted in a retrograde manner to post-transcriptionally regulate POR protein levels of two key transcription factors, PIF3 and HY5, in *ccr2* and *ccr2 det1-154* backgrounds (Figure 7). Of particular interest is how the abundance all three proteins was reverted in *ccr2 det1-154* to expected levels for *det1* mutants by treatment with D15. (Figure 7). Previous research has shown that *hy5*, *pif3* and *pifq* dark grown seedlings all contain etioplasts with PLBs (Chang et al., 2008; Stephenson et al., 2009; Martin et al., 2016). Consequently, we deduce that the lack of a PLB in *ccr2* is neither a consequence of apocarotenoid regulation of PIF3 or HY5, nor a lack of POR. An alternative hypothesis proposed in our model (Figure 8B) depicts how the apocarotenoid signal and DET1 may regulate an unknown factor required for PLB formation that is independent of POR abundance.

### An apocarotenoid signal regulates skotomorphogenesis and plastid biogenesis in parallel to DET1

DET1 is required for *cis*-carotene biosynthesis in wild type tissues, as *det1* mutants accumulate phytoene, phytofluene and tri-*cis*-ζ-carotene (Supplemental Table 7). c*is*-carotenes will hyper-accumulate in etiolated cotyledons and younger leaf tissues exposed to an extended dark period when CRTISO activity becomes rate-limited, such as in the absence of SDG8, which is required for permissive expression of *CRTISO* in the shoot meristem (Cazzonelli et al., 2009b; Cazzonelli et al., 2009a; Cazzonelli et al., 2010)(Figure 2). *SDG8* transcript levels are developmentally regulated, increasing from low basal levels after germination and declining during the dark phase of the diurnal cycle (Kim et al., 2005). Therefore, the accumulation of *cis*-carotenes and the apocarotenoid signal that regulates plastid biogenesis can be finely tuned with epigenetic and chromatin modifying processes that control development.

Herein we revealed how *ccr2* and *det1* oppositely regulate the chlorophyll biosynthetic enzyme, POR, at post-transcriptional and transcriptional levels, respectively, to control PLB formation (Figure 8). There are relatively few mutants published to date that do not produce a PLB in dark grown tissues and all, except *ccr2*, are due to reduced levels of PORA and/or PChlide. Arabidopsis mutants like *det1-1* and *cop1* mutants have less photoactive PChlide-F655 and higher total PChlide levels due to a reduction in POR that block PLB formation. Like *det1-1*, *det1-154* exhibits all the same phenotypes and indeed D15 treatment of *ccr2 det1-154* blocked PLB formation (Chory et al., 1989)(Supplementary Figure 3)(Table 1). Although, etioplasts in *ccr2* dark grown cotyledons do not make a PLB, they have an abundance of POR protein and total PChlide levels are similar to wild type (Figure 6,7). Therefore, *ccr2* and *det1* control PLB formation via independent signalling pathways. The *cis*-carotene derived cleavage product acts independent of *det1* to post-transcriptionally up-regulate POR protein levels and enhance PChlide thereby enabling PLB formation and etioplasts to chloroplast differentiation following de-etiolation and the normal greening of cotyledons exposed to continuous light.

*DET1* encodes a nuclear protein acting downstream from the phytochrome photoreceptors to regulate light-driven seedling development and *PhANG* expression (Schroeder et al., 2002). DET1 interacts with COP1 and the chromatin regulator DDB1, to limit the access of transcription factors to promoters and negatively regulate the expression of hundreds of genes via chromatin interactions (Schroeder et al., 2002; Lau and Deng, 2012). Light stimulates photomorphogenesis and the rapid down-regulation of *DET1* leading to a lower PIF3:HY5 protein ratio and the up-regulation of *PhANG* expression. Genetic mutations in *cop1* and *det*1 also lower the PIF3:HY5 ratio thereby activating *PhANG* expression (Benvenuto et al., 2002) (Osterlund et al., 2000). Consistent with these findings, *ccr2 det1-154* etiolated seedlings treated with D15 displayed elevated HY5 and almost negligible PIF3 protein levels, contrasting opposite to that of *ccr2*, revealing that the *cis*-carotene derived cleavage metabolite post-transcriptionally antagonises DET1 regulation of HY5 and PIF3 (Figure 8). This raised a question as to whether the *ccr2*-derived cleavage product could directly regulate DET1? This is unlikely for several reasons. First, *ccr2* and *ccr2 ziso-155* displayed closed cotyledons, an apical hook and normal hypocotyl length revealing that the *cis*-carotene derived cleavage metabolite does not activate photomorphogenesis (Table 1). Second, DET1 protein levels were relatively unchanged in WT, *ccr2* and *ccr2 det1-154*, regardless of D15 chemical inhibition. In conclusion, we deduce that the apocarotenoid signal acts in parallel with DET1 to regulate POR, PIF3 and HY5 protein accumulation and thus regulate etioplast development during skotomorphogenesis and chloroplast development under extended periods of darkness.

## METHODS

### Mutants used in this study

All germplasms are in the *Arabidopsis thaliana* ecotype Columbia (Col-0) background except where otherwise indicated. Germplasm used in this study include; *ziso*#11C (*zic1-3*: Salk_136385), *ziso*#12D (*zic1-6*; Salk_057915C), *ziso*#13A (*zic1-4*; CS859876), *ccr2.1/crtiso* (Park et al., 2002), *ccr1.1*/*sdg8* (Cazzonelli et al., 2009b), *lut2-1* (Pogson et al., 1996), *ccd1-1* (SAIL_390_C01), *ccd4* (Salk_097984c), *max3-9/ccd7* (Stirnberg et al., 2002), *max4-1/ccd8* (Sorefan et al., 2003), *aba1-3* (Koornneef et al., 1982), *det1-1* (CS6158), *ccr2 det1-154*, and *det1-154*.

### Plant growth conditions and treatments

For soil grown plants, seeds were sown on DEBCO seed raising mixture and stratified for 3 d at 4 °C in the dark, prior to transferring to an environmentally controlled growth chamber set to 21 °C and illuminated by approximately 120 µmol.m^−2^.sec^−1^ of fluorescent lighting. Unless otherwise stated, plants were grown in a 16-h photoperiod. Photoperiod shift assays were performed by shifting 2-3 week old plants grown under a 16-h photoperiod to an 8-h photoperiod for one week and newly emerged immature leaves were scored as displaying either a yellow leaf (YL) or green leaf (GL) phenotype, reflecting either impaired or normal plastid development respectively.

For media grown seedlings, Arabidopsis seeds were sterilized for 3 h under chlorine gas in a sealed container, followed by washing seeds once with 70% ethanol and three times with sterilized water. Seeds were sown onto Murashige and Skoog (MS) media (Caisson Labs; MSP01) containing 0.5% phytagel (Sigma) and half-strength of Gamborg’s vitamin solution 1000X (Sigma Aldrich) followed by stratification for 2 d (4 °C in dark) to synchronise germination. Inhibition of carotenoid cleavage dioxygenase (CCD) enzyme activity was achieved by adding D15 (aryl-C3N hydroxamic acid) dissolved in ethanol to a final optimal concentration of 100 μM as previously described (Van Norman et al., 2014). Etiolation experiments involved growing seedlings in the dark at 21°C for 7 d and harvesting tissue under a dim green LED light. For de-etiolation and greening experiments, Arabidopsis seeds were stratified for 2 d and germinated in the dark at 21°C for 4 d. Seedlings were then exposed to constant light (~80 µmol.m^−2^.sec^−1^, metal-halide lamp) for 72 h at 21 °C. Cotyledon tissues were harvested at 24-h intervals for chlorophyll quantification.

### Plasmid construction

pEARLEY::ZISO-OE and pEARLEY::DET1-OE binary vectors were designed to overexpress ZISO and DET1 cDNA fragments, respectively. Both genes were regulated by the constitutive CaMV35S promoter. Full length cDNA coding regions were chemically synthesised (Thermo Fisher Scientific) and cloned into the intermediate vector pDONR221. Next, using gateway homologous recombination, the cDNA fragments were cloned into pEarleyGate100 vector as per Gateway^®^ Technology manufacturer’s instructions (Thermo Fisher Scientific). Vector construction was confirmed by restriction digestion and sanger sequencing.

### Generation of transgenic plants

The *ccr2 ziso-155* and *ccr2 det1-154* EMS generated mutant lines were transformed by dipping Arabidopsis flowers with Agrobacteria harbouring pEARLEY::ZISO-OE or pEARLEY::DET1-OE binary vectors to generate *ccr2 ziso-155*::ZISO-OE and *ccr2 det1^154^*::DET1-OE transgenic lines, respectively. At least 10 independent transgenic lines were generated by spraying seedlings grown on soil with 50 mg/L of glufosinate-ammonium salt (Basta herbicide).

### Chlorophyll pigment quantification

Total chlorophyll was measured as described previously (Porra et al., 1989) with minor modifications. Briefly, 20 seedlings from each sample were frozen and ground to fine powder using a TissueLyser (Qiagen). Homogenised tissue was rigorously suspended in 300 µL of extraction buffer (80% acetone and 2.5mM NaH_2_PO_4_, pH 7.4), incubated at 4°C in dark for 15 min and centrifuged at 20,000 g for 10 min. Two hundred and fifty microliters of supernatant was transferred to a NUNC 96-well plate (Thermo Fisher Scientific) and measurements of A647, A664 and A750 were obtained using an iMark Microplate Absorbance Reader (Thermo Fisher Scientific). Total chlorophyll in each extract was determined using the following equation modified from (Porra, 2002): (Chl a + Chl b) (µg) = (17.76 × (A647-A750) + 7.34 × (A664-A750)) × 0.895 × 0.25.

### Carotenoid pigment analysis

Pigment extraction and HPLC-based separation was performed as previously described (Cuttriss et al., 2007; Dhami et al., 2018). Reverse phase HPLC (Agilent 1200 Series) was performed using either the GraceSmart-C18 (4-µm, 4.6 × 250-mm column; Alltech) or Allsphere-C18 (OD2 Column 5-µm, 4.6 × 250; Grace Davison) and/or YMC-C30 (250 × 4.6mm, S-5µm) columns. The C18 columns were used to quantify β-carotene, xanthophylls and generate *cis*-carotene chromatograms, while the C30 column improved *cis*-carotene separation and absolute quantification. Carotenoids and chlorophylls were identified based upon retention time relative to known standards and their light emission absorbance spectra at 440 nm (chlorophyll, β-carotene, xanthophylls, pro-neurosporene, tetra-cis-lycopene), 400 nm (-carotenes), 340 nm (phytofluene) and 286 nm (phytoene). Quantification of xanthophyll pigments was performed as previously described (Pogson et al., 1996). Quantification of *cis*-carotenes was performed by using their molar extinction coefficient and molecular weight to derive peak area in terms of micrograms (μg) per gram fresh weight (gfw) as previously described (Britton, 1995).

### Transmission Electron Microscopy (TEM)

Cotyledons from 5-d-old etiolated seedlings were harvested in dim-green safe light and fixed overnight in primary fixation buffer (2.5% Glutaraldehyde and 4% paraformaldehyde in 0.1 M phosphate buffer pH 7.2) under vacuum, post-fixed in 1% osmium tetroxide for 1 h, followed by an ethanol series: 50%, 70%, 80%, 90%, 95% and 3 × 100% for 10 min each. After dehydration, samples were incubated in epon araldite (resin): ethanol at 1 : 2, 1 : 1 and 2:1 for 30 min each, then 3 times in 100% resin for 2 h. Samples were then transferred to fresh resin and hardened under nitrogen air at 60 °C for 2 d, followed by sectioning of samples using Leica EM UC7 ultramicrotome (Wetzlar). Sections were placed on copper grids, stained with 5% uranyl acetate, washed thoroughly with distilled water, dried, and imaged with H7100FA transmission electron microscope (Hitachi) at 100 kV. For each of the dark-grown seedling samples, prolamellar bodies were counted from 12 fields on 3 grids, and data analysed using two-way ANOVA with post-hoc Tukey HSD.

### DNA-seq Library Construction, Sequencing and Bioinformatics Identification of SNPs

Genomic DNA (gDNA) was extracted using the DNeasy Plant Mini Kit (Qiagen). One microgram of gDNA was sheared using the M220 Focused-Ultrasonicator (Covaris) and libraries were prepared using NEBNext^®^ Ultra™ DNA Library Prep Kit (New England Biolabs) followed by size selection (~320 bp) using AMPure XP Beads (Beckman Coulter). Paired-end sequencing was performed using the Illumina HiSEQ1500. After sequencing, the raw reads were assessed for quality using the FastQC software (http://www.bioinformatics.babraham.ac.uk/projects/fastqc/), and subjected to trimming of illumina adapters and filtering of low quality reads with AdapterRemoval programme (Lindgreen, 2012). The reads were mapped to the *Arabidopsis thaliana* (TAIR9) genome with BWA mapper (Li and Durbin, 2009). The resultant BWA alignment files were converted to sorted bam files using the samtools v0.1.18 package (Li et al., 2009) and were used as input for the subsequent SNP calling analyses. The SNPs were called and analysed further on both the parent and mutant lines using NGM pipeline (Austin et al., 2011) and SHOREmap (Schneeberger et al., 2009). For the NGM pipeline, SNPs were called using samtools (v0.1.16) as instructed and processed into ‘.emap’ files using a script provided on the NGM website. The .emap files were uploaded to the NGM web-portal to assess SNPs with associated discordant chastity values. To identify mutant specific SNPs, SNPs from parental lines were filtered out and EMS-induced homozygous SNPs were defined based on the discordant chastity metric. For SHOREmap, the SHORE software (Ossowski et al., 2008) was used to align the reads (implementing BWA) and call the SNPs (Hartwig et al., 2012). SHOREmap backcross was then implemented to calculate mutant allele frequencies, filter out parent SNPs and define the EMS mutational changes. Where appropriate, custom scripts were used to identify mutant specific EMS SNPs, filter out parent SNPs and annotate the region of interest. The SNPs and InDels were localized based on the annotation of gene models provided by TAIR database (http://www.arabidopsis.org/). The polymorphisms in the gene region and other genome regions were annotated as genic and intergenic, respectively. The genic polymorphisms were classified as CDS (coding sequences), UTR (untranslated regions), introns and splice site junctions according to their localization. SNPs in the CDS were further separated into synonymous and non-synonymous amino substitution. The GO/PFAM annotation data were further used to functionally annotate each gene.

### RNA-seq Library Construction, Sequencing and Differential Gene Expression Analysis

Total RNA was extracted from Arabidopsis leaf tissues grown under an 8-h photoperiod or cotyledons from etiolated seedlings grown in dark for 7 d by TRIzol (Thermo Fisher Scientific) followed by DNase treatment at 37 °C for 30 min. RNA was recovered using x1.8 Agencourt RNAClean XP magnetic beads (Beckman Coulter). RNA (1 µg) libraries were constructed using 1llumina TruSeq Stranded mRNA Library Prep Kit (ROCHE) followed by bead size selection (~280 bp) using AMPure XP Beads and libraries sequenced using the Illumina HiSEQ2000. Fifteen million reads were obtained from sequencing each library and 21365 to 23840 mRNA transcripts were identified. Quality control was performed with FASTQC v.0.11.2. Adapters were removed using scythe v.0.991 (flags -p 0.01 for the prior), reads trimmed with sickle v.1.33 (flags q 20; quality threshold and −l 20 for minimum read length after trimming) and aligned to the Arabidopsis genome (TAIR10) using the subjunc v.1.4.6 aligner (-u and -H flags to report reads with a single, unambiguous mapping location) (Liao et al., 2014). The number of reads mapping per gene were summarised using feature Counts (v.1.4.6 with flags -s 2, -P and -c) to map reverse stranded and discard read pairs mapping to different chromosomes (Liao et al., 2014). Statistical testing for relative gene expression was performed in R using edgeR v.3.4.2 (Robinson and Smyth, 2007, 2008; Robinson et al., 2010; Robinson and Oshlack, 2010; McCarthy et al., 2012), Voom (Law et al., 2014) in the limma package 3.20.1 (Smyth, 2004, 2005). Transcripts were considered differentially expressed when a fold change > 2 and FDR adjusted *p* < 0.05. The bioinformatics analysis pipeline from fastq to summarised counts per gene is available at https://github.com/pedrocrisp/NGS-pipelines. RNAseq data sets was deposited into a permanent public repository with open access (https://www.ncbi.nlm.nih.gov/sra/PRJNA498324).

### Protein extraction and western blot analysis

For protein extraction, fifty to one hundred milligrams of etiolated Arabidopsis cotyledons (7-d-old) were harvested under dim-green safe light and ground to fine powder. Total protein was extracted using a TCA-acetone protocol (Mechin et al., 2007) with minor modification and pellets were resuspended in 100 μL – 200 μL solubilization buffer. The concentration of total protein was measured using Bradford reagent (Bio-Rad) and adjusted to 2 μg/μL. A serial dilution was used to determine western blot sensitivity for each antibody and determine the optimal concentration for quantification. Five micrograms of total protein run on a gel was transferred to a PVDF membrane (Bio-Rad) and incubated with anti-POR polyclonal antibody (Agrisera Antibodies AS05067, 1:2000), anti-PIF3 polyclonal antibody (Agrisera Antibodies, 1:2000) or anti-HY5 antibody (Agrisera Antibodies AS05067, 1:1000) for 2 h. To examine DET1 protein levels, 10 μg of total protein was loaded to the gel and anti-DET1 polyclonal antibody (Agrisera Antibodies AS153082) was used at a 1:1000 dilution. Membranes were washed and incubated with HRP-conjugated Goat anti-Rabbit IgG (Agrisera Antibodies, 1:2500) for 90 min. Membranes were re-probed using anti-Actin polyclonal antibody (Agrisera Antibodies AS132640, 1:3000) and HRP-conjugated Goat anti-Rabbit IgG (Agrisera Antibodies, 1:2500) for internal protein normalisation.

### Protochlorophyllide Quantification

Protochlorophyllides (Pchlides) were extracted and measured using published methods (Kolossov and Rebeiz, 2003) with modifications. Around 100 mg of etiolated Arabidopsis seedlings (7-d-old) were harvested under dim-green safe light, frozen and ground to fine powder. Two milliliters of 80% ice-cold acetone was added to each sample and the mixture was briefly homogenized. After centrifugation at 18,000 g for 10 min at 1 °C, supernatant was split to 2 × 1 mL for Pchlides and protein extraction. Fully esterified tetrapyrroles were extracted from the acetone extracts with equal volume followed by 1/3 volume of hexane. Pchlides remained in the hexane-extracted acetone residue were used for fluorescence measurement with a TECAN M1000PRO plate reader (Tecan Group) and net fluorescence were determined as previously described (Rebeiz et al., 1975). Protein extraction was performed using 80% acetone and 10% TCA; protein concentration was used to normalize the net fluorescence of Pchlides.

### Real-Time PCR Analysis

The total RNA was extracted using Spectrum™ Plant Total RNA kit as per manufacturer’s protocol (Sigma-Aldrich). The qRT-PCR was performed with mixture of 2 μL of primer mix (2 μM from each F & R primer), 1 μL 1/10 diluted cDNA template, 5 μL LightCycler 480 SYBR Green I Master mix and distilled water up to a total volume of 10 μL. Relative transcript abundance was quantified using LightCycler 480 as per instructions (Roche). For each sample, three technical replicates for each of three biological replicates were tested. The relative gene expression levels were calculated by using relative quantification (Target Eff Ct(Wt-target) / Reference Eff Ct(Wt-target)) and fit point analysis (Pfaffl, 2001). Protein Phosphatase 2A (At1g13320) was used as housekeeper reference control for all experiments (Czechowski et al., 2005). All primer sequences are listed in Supplemental Table 8. Statistical analysis was performed using Two-Way ANOVA.

## SUPPORTING INFORMATION

**Supplemental Figure 1.**
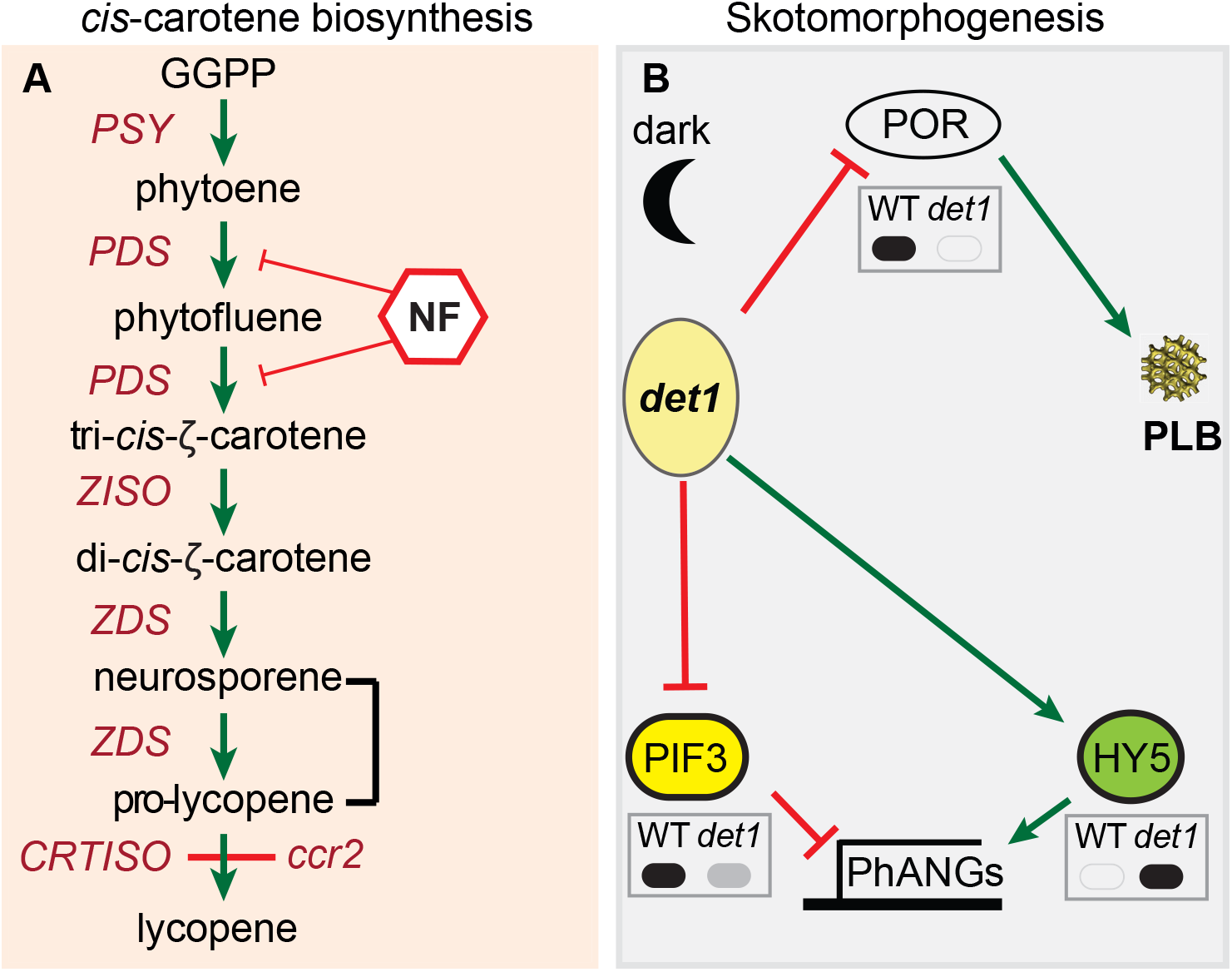
A model for *cis*-carotene biosynthesis and regulation of PLB formation during skotomorphogenesis. A) A pathway for *cis*-carotene biosynthesis. The carotenoid isomerase mutant (*ccr2*) accumulates *cis*-carotenes in etiolated seedlings. Norflurazon (NF) inhibits *cis*-carotene accumulation. B) Control of prolamellar body (PLB) formation and protein levels during skotomorphogenesis. DET1 acts as a repressor of photomorphogenesis in etiolated tissues to maintain high protein levels of PIF3, which reduce *PhANG* expression. Upon de-etiolation, DET1 and PIF3 protein levels decline and *det1* mutants accumulate HY5 protein, which promotes the expression of *PhANGs*. *det1* mutants do not accumulate PORA proteins and do not form a PLB in etioplasts. Grey insert boxes digitally represent published western protein blots for PORA (Lebedev et al., 1995), PIF3 (Dong et al., 2014) and HY5 (Osterlund et al., 2000) in WT and *det1* mutant genotypes. Solid black and grey fills represents high and low protein expression, respectively. Green arrows and red lines represent positive and negative regulation, respectively. Abbreviations: PSY, phytoene synthase; PDS, phytoene desaturase, ZDS, ζ-carotene desaturase; Z1SO, ζ-carotene isomerase; CRTISO, carotenoid isomerase

**Supplemental Figure 2.**
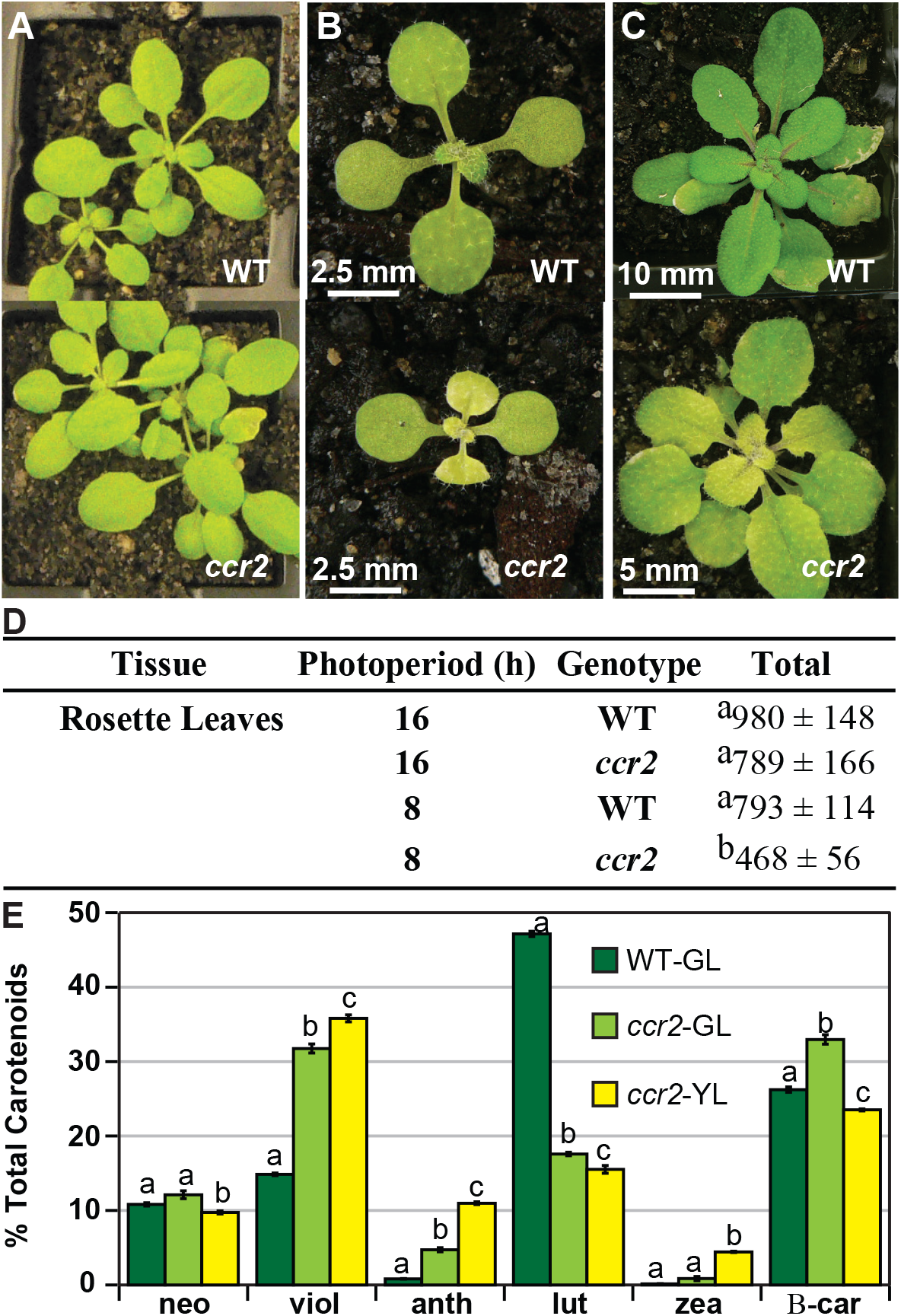
A shorter photoperiod promotes leaf variegation affecting chlorophyll levels and carotenoid composition in *ccr2*. **(A)** WT and *ccr2* plants were grown under a lower intensity of light (50 μmol m^−2^ s^−1^) and representative images taken 14 DAG. **(B)** and **(C)** WT and *ccr2* plants were grown under a very short 8 h photoperiod and representative images taken after 14 (B) and 21 (C) days of growth. **(D)** Chlorophyll content in immature leaves that recently emerged from WT and ccr2 rosettes 14 DAG. Values represent the average and standard deviations of total chlorophyll content (μg/gfw) from a single leaf sector (n=2-7 plants). Lettering denotes significance (ANOVA, *p* < 0.05). **(E)** Percentage carotenoid composition (relative to total) in green (WT and *ccr2*) and yellow (*ccr2*) virescent leaves developed one week after a 16 h to 8 h photoperiod shift. Values represent average and standard error of means are displayed (n=5, single leaf from 5 plants). Lettering denotes significance (paired *t*-test; *p* < 0.05). Neoxanthin (neo), violaxanthin (viol), antheraxanthin (anth), lutein (lutein), zeaxanthin (zea), β-car (β-carotene), Green Leaf (GL), Yellow Leaf (YL).

**Supplemental Figure 3.**
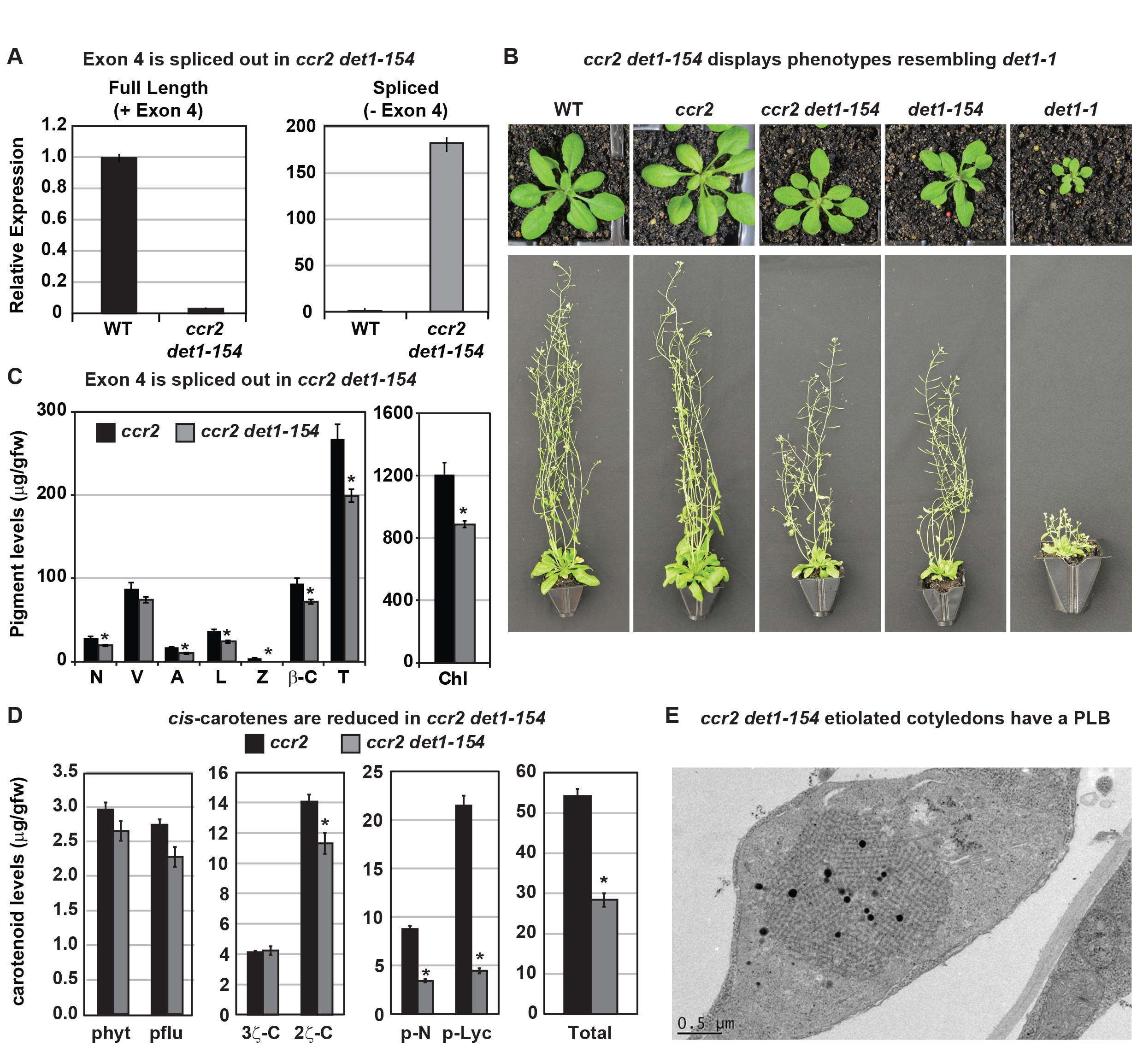
*det1-154* has alternative splicing and reduced pigments, *cis*-carotenes and restored PLB formation in *ccr2*. **(A)** qRT-PCR confirms alternative splicing of exon 4 in *ccr2 det1-154* leaf tissues. Primers were designed to quantify the full length (+ Exon 4; spanning exons 3-4 and 4-5 junctions) and the spliced (- Exon 4: spanning exon 3-5 and 6-7 junctions) *DET1*-*154* mRNA transcript levels in WT and *ccr2 det1-154* leaf tissues, respectively. Standard error bars are shown (n=4). **(B)** *ccr2 det1-154* displays phenotypes resembling *det1-1*, including a small rosette, shorter floral architecture and partially sterility in comparison to WT and *ccr2*. **(C)** *ccr2 det1-154* shows reduced pigment levels compared to *ccr2*. Neoxanthin (N); violaxanthin (V); antheraxanthin (A), lutein (L), β-carotene (β-C), total carotenoids (T) and total chlorophylls (Chl) were quantified at a 440nm. Mean values are displayed and error bars denote standard error (n=3). Star denotes significance (ANOVA, p < 0.05). Data is representative of multiple experiments. **(D)** *det1-154* reduces *cis*-carotene content in *ccr2*. phytoene (phyt), phytofluene (pflu), tri-*cis*-ζ-carotene (3ζ-C), di-*cis*-ζ-carotene (2ζ-C), pro-neurosporene (p-N), tetra-*cis*-lycopene (p-lyc) and total *cis*-carotenes were quantified at absorption wavelengths providing maximum detection. Star denotes significance (ANOVA, *p* < 0.05). Data is representative of two independent experiments and error bars show standard error (n=4). **(D)** Transmission electron micrographs of a representative etioplast from 5-d-old dark grown cotyledons showing a well-developed PLB in *ccr2 det1-154*.

**Supplemental Figure 4.**
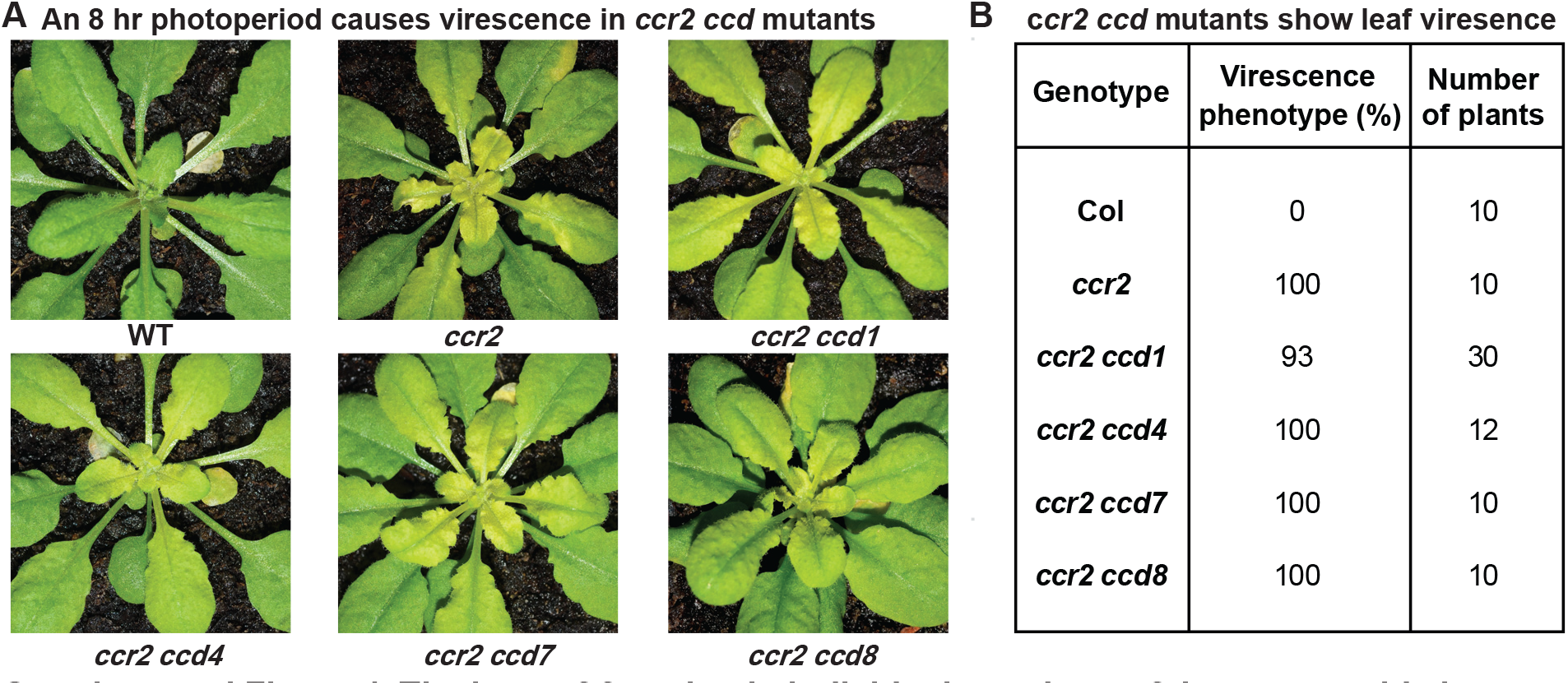
The loss-of-function in individual members of the *carotenoid cleavage dioxygenase* gene family cannot restore plastid development in *ccr2* rosettes. Three-week-old WT, *ccr2*, *ccr2 ccd1, ccr2 ccd4, ccr2 ccd7*, and *ccr2 ccd8* (F_3_ homozygous double mutant lines) plants were shifted from a 16-h to 8-h photoperiod until newly formed leaves in the *ccr2* rosette displayed a virescent leaf phenotype. **(A)** Representative images of plants showing newly developed leaves in the rosette. **(B)** Quantification of leaf variegation in individual rosettes from *ccr2 ccd* double mutants. Data is representative of multiple independent experiments. Statistical analysis by ANOVA with post-hoc Tukey test showed no significant difference in the number of *ccr2* and *ccr2 ccd* plants displaying a virescent phenotype.

**Supplemental Table 1.** Immature *ccr2* tissues have an altered *cis*-carotene and xanthophyll composition.

| Genotype | Tissue | Age | Percentage xanthophyll composition |  |  |  |  |  | Relative ratio |  |
| --- | --- | --- | --- | --- | --- | --- | --- | --- | --- | --- |
| | | | lutein | β-c | zea | anth | viol | neo | phyt | pflu |
| WT | Rosette leaf | yng | 50 | 26 | 0.0 | 0.5 | 10 | 13 | 0.00 | 0.00 |
|  | Rosette leaf | old | 48 | 27 | 0.1 | 0.6 | 11 | 13 | 0.00 | 0.00 |
|  | Floral bud | yng | 50 | 24 | 0.0 | 0.6 | 15 | 10 | 0.00 | 0.00 |
|  | Floral bud | old | 51 | 23 | 0.0 | 0.4 | 16 | 9 | 0.00 | 0.00 |
|  | Maximum SD |  | 1 | 1 | 0.3 | 0.4 | 0 | 2 | ND | ND |
| *ccr2* | Rosette leaf | yng | 11 | 33 | 1.4 | 5.5 | 36 | 13 | 0.37 | 0.29 |
|  | Rosette leaf | old | 20 | 36 | 0.6 | 3.4 | 28 | 12 | 0.02 | 0.00 |
|  | Floral bud | yng | 12 | 34 | 1.3 | 4.2 | 38 | 11 | 1.55 | 0.59 |
|  | Floral bud | old | 17 | 34 | 1.0 | 3.5 | 33 | 12 | 0.76 | 0.38 |
|  | Maximum SD |  | 1.2 | 0.6 | 0.2 | 0.7 | 0.7 | 0.3 | nd | nd |

Percentage of individual carotenoid levels relative to the total carotenoid content in different tissues from plants exposed to a 16 h photoperiod. Ratios of phytoene and phytofluene are relative to β-carotene. Data represent the average and maximum standard deviation (SD) for 2 biological replicates. Similar results were observed in independent experiments. Greyed highlighted values represent significant (t-test $p < 0.05$ ) differences in immature younger (yng) relative to mature older (old) tissues. $\beta$ -c; β-carotene, zea; zeaxanthin, anth; antheraxanthin, viol; violaxanthin, neo; neoxanthin, phyt; phytoene, pflu; phytofluene. nd; not determined.

**Supplemental Table 2.** D15 and *ziso* restore PLB formation in *ccr2* etiolated cotyledons.

| Germplasm | Treatment | Number of etioplasts containing PLBs |  |  |  | Tukey Groups |
| --- | --- | --- | --- | --- | --- | --- |
|  |  | Total | PLB | Ratio (%) | SE (%) |  |
| WT | H<sub>2</sub>O | 90 | 90 | 100 | 0.0 | c |
| WT | EtOH | 64 | 64 | 100 | 0.0 | c |
| WT | D15 | 48 | 48 | 100 | 0.0 | c |
| *ccr2* | H<sub>2</sub>O | 63 | 0 | 0 | 0.0 | a |
| *ccr2* | EtOH | 72 | 0 | 0 | 0.0 | a |
| *ccr2* | D15 | 71 | 61 | 85 | 2.5 | b |
| *ziso1-4* | H<sub>2</sub>O | 73 | 48 | 66 | 2.1 | d |
| *ziso1-4* | D15 | 68 | 45 | 66 | 2.3 | d |
| *ccr2 ziso1-4* | H<sub>2</sub>O | 63 | 59 | 94 | 5.5 | c |
| *ccr2 ziso-155* | H<sub>2</sub>O | 79 | 76 | 95 | 5.0 | c |

PLB formation was examined in WT, *ccr2*, *ziso*, *ccr2 ziso* and *ccr2 ziso-155* cotyledons 7 DAG in the dark. D15 (CCD inhibitor), EtOH (control solvent for dissolving D15) and/or H<sub>2</sub>O was added to the growth media treatments.

**Supplemental Table 3.** Transcriptomic analysis of WT, *ccr2* and *ccr2 ziso-155* etiolated tissues.

**Supplemental Table 4.** Transcriptome analysis of WT, *ccr2* and *ccr2 ziso-155* immature leaf tissues

**Supplemental Table 5.** Significantly expressed genes regulated in *ccr2* and contra-regulated *ccr2 ziso-155* that are common to both etiolated and immature leaf tissues.

**Supplemental Table 6.** Contra-regulated differential gene expression in etiolated seedlings and young leaves of *ccr2 ziso-155.*

| GENE ID | GENE | Protein Encoding Description | Etiolated Seedlings |  | Young Leaves |  | *det1.1* | NF |
| --- | --- | --- | --- | --- | --- | --- | --- | --- |
|  |  |  | *ccr2* | *ziso-155* | *ccr2* | *ziso-155* |  |  |
| At1g09530 | PIF3 | Transcription factor interacts with photoreceptors and negatively regulates signalling | 30 | 0.1 | 220 | 0.1 | ↓ | NS |
| At4g10180 | DET1/FUS2 | Encodes a nuclear-localized protein that acts as a repressor of photomorphogenesis | 5.1 | 0.1 | 5.9 | 0.2 | NS | NS |
| At3g19390 |  | Granulin repeat cysteine protease family protein | 4.4 | NS | 6.8 | NS | NS | NS |
| At5g13210 |  | Unknown conserved expressed protein | 3.8 | NS | 0.4 | NS | ↑ | NS |
| At3g45730 |  | Unknown expressed protein | 2.8 | NS | 2.4 | NS | NS | 10.6 |
| At5g43500 | ATARP9 | Encodes an expressed protein similar to actin-related proteins | 2.4 | NS | 2.2 | NS | NS | NS |
| At5g48240 |  | Unknown expressed protein | 2.1 | NS | 2.2 | NS | NS | NS |
| At2g32950 | COP1/FUS3 | Repressor of photomorphogenesis and induces skotomorphogenesis in the dark | 2.0 | 0.0 | 8.9 | 0.1 | ↑ | NS |
| At5g11260 | HY5 | Transcription factor negatively regulated by COP1 and mutant shows ABA resistant phenotypes | 0.5 | 8.1 | 0.3 | 8.4 | NS | 2.8 |
| At4g02770 | PSAD1 | Expressed protein with similarity to photosystem I reaction center subunit II | 0.5 | NS | 0.5 | NS | ↑ | 0.15 |
| At3g17070 |  | Peroxidase family expressed protein | 0.5 | NS | 0.5 | NS | NS | NS |
| At2g31751 |  | Potential natural antisense gene, expressed protein | 0.4 | NS | 0.5 | NS | NS | NS |
| At4g15560 | DXS/CLA1 | 1-deoxyxylulose 5-phosphate synthase activity involved in the MEP pathway | 0.3 | 4.2 | 0.1 | 16.2 | NS | 0.42 |
| At4g34350 | ISPH/CLB6 | 4-hydroxy-3-methylbut-2-enyl diphosphate reductase involved in the MEP pathway | 0.3 | 9.4 | 0.2 | 11 | ↑ | NS |
| At1g24510 | TCP-1 | T-complex expressed protein 1 epsilon subunit | 0.3 | 12.0 | 0.1 | 7.9 | NS | NS |
| At3g59010 | PME35 | Pectin methylesterase that regulates the cell wall mechanical strength | 0.2 | NS | 0.4 | NS | ↓ | NS |
| At1g29930 | CAB1/LHCB1.3 | Subunit of light-harvesting complex II (LHCII), which absorbs light and transfers energy to the photosynthetic reaction center | 0.2 | 13 | 0.2 | 11 | NS | NS |
| At2g05070 | LHCB2.2 | Light-harvesting chlorophyll a/b-binding (LHC) protein that constitute the antenna system of the photosynthetic apparatus. | 0.2 | NS | 0.2 | NS | ↑ | NS |
| At5g13630 | GUN5/CHLH | Magnesium chelatase involved in plastid-to-nucleus signal transduction. | 0.2 | 17 | 0.2 | 20 | ↑ | 0.33 |
| At1g67090 | RBCS1a | Member of the Rubisco small subunit (RBCS) multigene family and functions to yield sufficient Rubisco content for leaf photosynthetic capacity. | 0.1 | 67 | 0.1 | 61 | NS | NS |

Notes: NS; not significant. Transcriptomic data; *det1.1* (Schroeder, 2001) and norflurazon (NF) (Koussevitzky, 2007).

**Supplemental Table 7.**
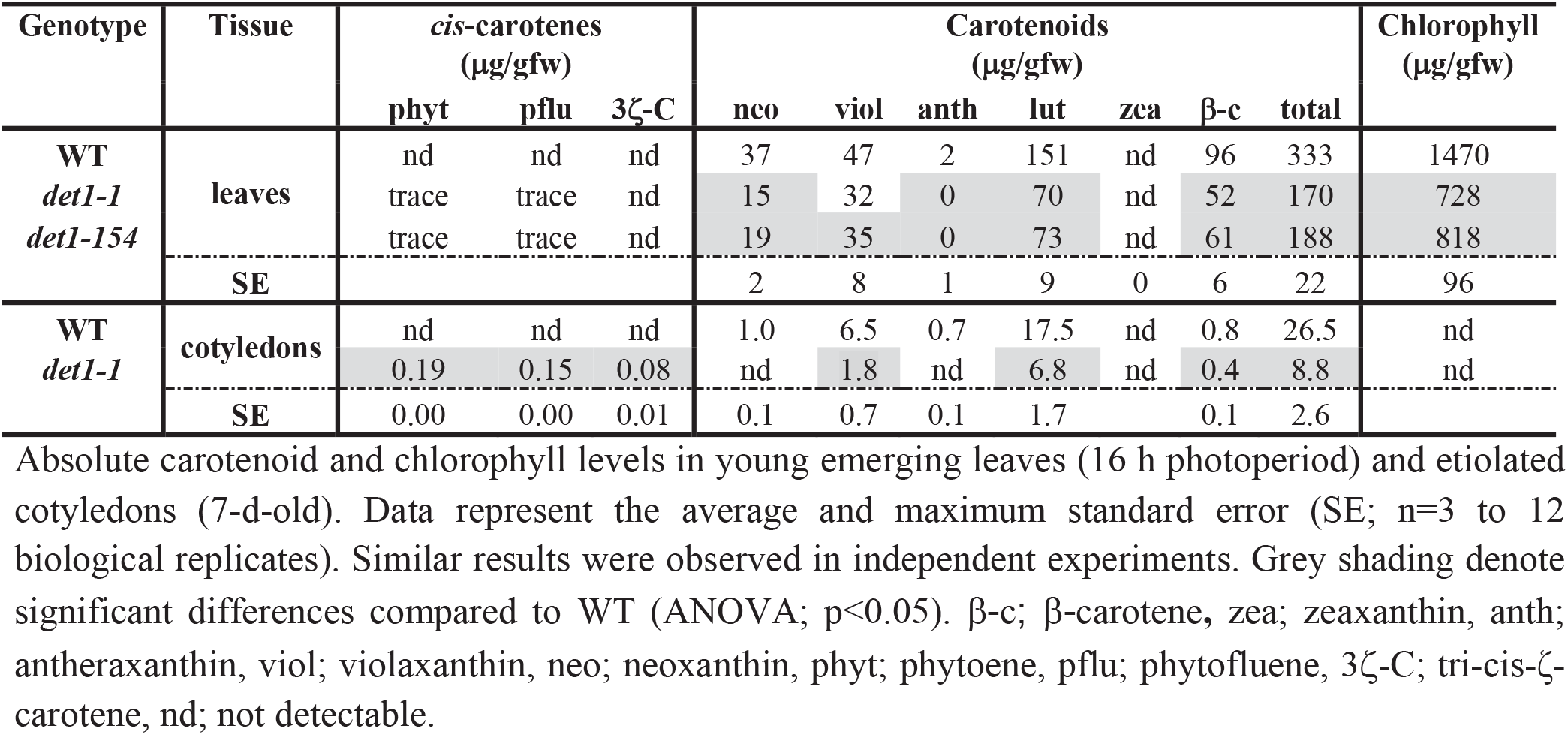
*det1* reduced carotenoids and caused *cis*-carotenes to accumulate in leaves and etiolated tissues.

**Supplemental Table 8.**
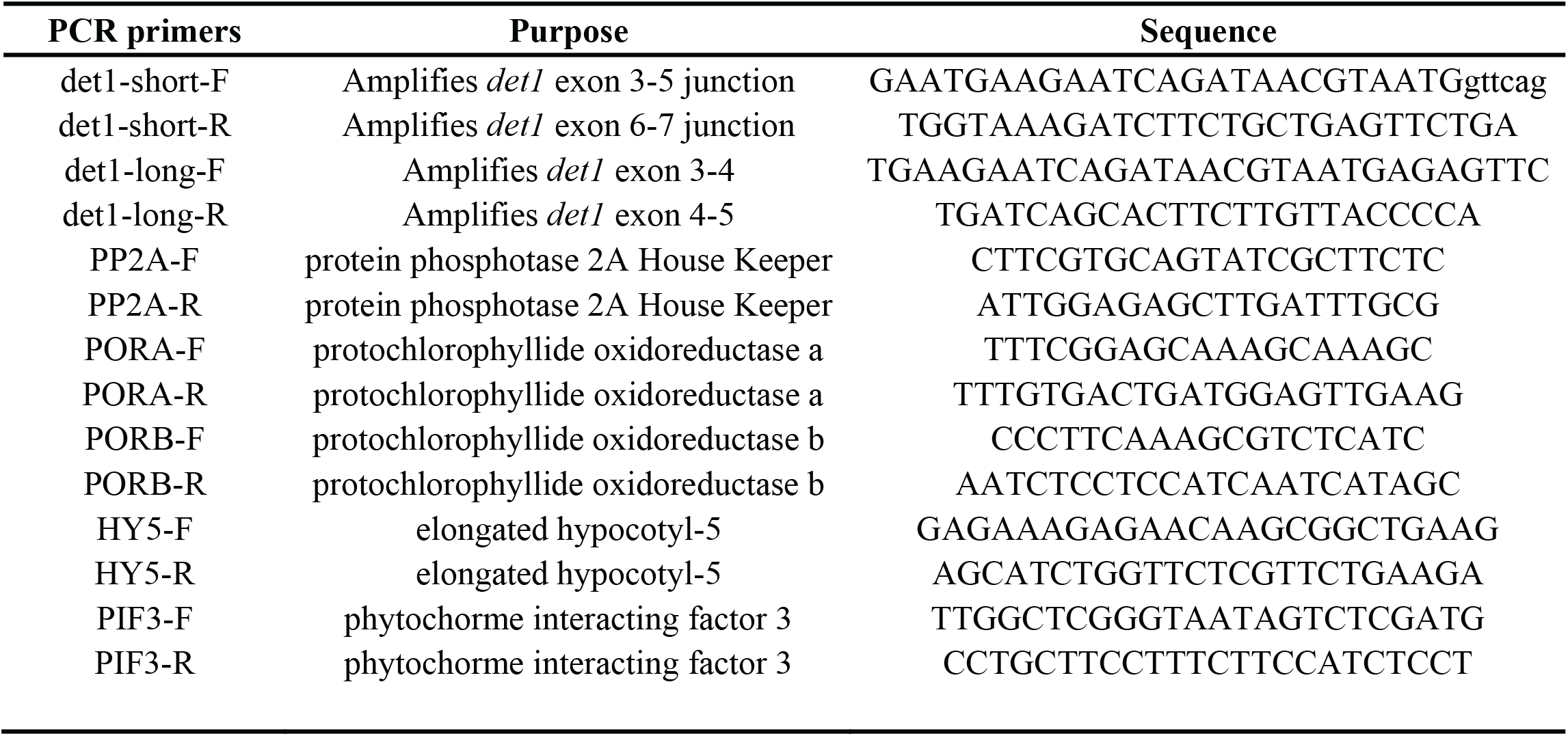
Primer sequences used for qRT-PCR and *ccr2 det154* characterisation

## ACKNOWLEDGEMENTS

We would like to especially thank Rishi Aryal for his technical assistance in confirming the splicing of *det1-154* and Peter Crisp for assisting XH with the bioinformatics analysis. Many thanks to Arun Yadav, Shelly Verma, William Walker, Michelle Nairn, Sam Perotti, Jacinta Watkins and Kai Chan for their assistance in maintaining plants, crossing mutant germplasm and performing HPLC. We thank Chris Mitchell for developing *cis*-carotene standards. We thank Philip Benfey for providing the D15 chemical inhibitor of CCD activity. Next generation sequencing was performed at the Biomolecular Resource Facility (ANU). This work was supported by Grant CE140100008 (BJP) and DP130102593 (CIC).

## AUTHOR CONTRIBUTIONS

CIC and BJP conceived ideas and designed research. CIC and XH wrote the manuscript. CIC and XH prepared figures and tables and performed the majority of experiments. YA contributed to Figures 5, 6, 7, 8. YA and ND contributed to Tables 1, S7 and Figure S3. JR produced Figure S3. MS contributed expertise in DNA bioinformatics analysis. JL contributed expertise in TEM. CIC and BJP supervised XH, JR and ND. CIC supervised YA. All authors contributed to editing the manuscript.

## Abbreviations

ccr: *carotenoid and chloroplast regulation*
rccr2: revertant of *ccr2*
DAG: days after germination
YL: yellow leaf
GL: green leaf
NF: norflurazon
ACS: apocarotenoid signal

